# *Dlk1-Dio3* Locus-Derived LncRNAs Perpetuate Postmitotic Motor Neuron Cell Fate and Subtype Identity

**DOI:** 10.1101/320135

**Authors:** Ya-Ping Yen, Wen-Fu Hsieh, Ya-Lin Lu, Ee Shan Liau, Ho-Chiang Hsu, Yen-Chung Chen, Ting-Chun Liu, Mien Chang, Joye Li, Shau-Ping Lin, Jui-Hung Hung, Jun-An Chen

## Abstract

The mammalian imprinted *Dlk1-Dio3* locus produces multiple long non-coding RNAs (lncRNAs) from the maternally inherited allele, including *Meg3* (i.e. *Gtl2*) in the mammalian genome. Although this locus has well-characterized functions in stem cell and tumor contexts, its role during neural development is unknown. By transcriptome profiling cell types at each stage of spinal cord development, we uncovered that lncRNAs expressed from the *Dlk1-Dio3* locus are predominantly and gradually enriched in rostral motor neurons (MNs). Mechanistically, *Meg3* and other *Dlk1-Dio3* locus-derived lncRNAs facilitate Jarid2-Ezh2 interactions. Loss of these lncRNAs compromises the H3K27me3 landscape, leading to aberrant expression of progenitor and caudal *Hox* genes in postmitotic MNs. Our data illustrate that these lncRNAs in the *Dlk1-Dio3* locus play a critical role in maintaining postmitotic MN cell fate by repressing progenitor genes and they shape MN subtype identity by regulating *Hox* genes, providing strong evidence of how lncRNAs function during embryonic development.

## INTRODUCTION

Investigations of the gene regulatory networks involved in cell type specification during embryonic development have been protein-centric for decades. However, given the prevalence of high-throughput sequencing analyses of mammalian genomes, it is now appreciated that non-coding RNAs (ncRNAs) account for at least 50~80 % of transcriptomes (Pauli et al, 2011; Rinn & Chang, 2012). Regulatory ncRNAs can be broadly classified based on their size (Mattick, 2009). Short RNA species (~20–30 nucleotides [nt]), such as microRNAs (miRNAs), have emerged as pivotal modulators of development and disease through mediation of translational repression or mRNA degradation (Esteller, 2011). Long non-coding RNAs (lncRNAs; > 200 nt) are gaining prominence for their roles in many cellular processes, from chromatin organization to gene expression regulation during embryonic development (Kung et al, 2013; Rinn & Chang, 2012; Rutenberg-Schoenberg et al, 2016). Thus, it is not surprising that lncRNAs were recently found to be associated with an array of diseases including cancers, as well as cardiovascular and neurological disorders (Briggs et al, 2015).

Accumulating evidence supports that lncRNAs can induce *cis*- and *trans-acting* gene silencing. For example, the lncRNA *Airn* directly represses the paternally-expressed *Igf2r* gene *in cis* for the maintenance of ESC differentiation (Pauler et al, 2005) and *Xist* lncRNA triggers *in cis* inactivation of the X chromosome (Lee, 2009). The human lncRNA *HOTAIR*, which is expressed from the caudal *HOXC* locus, acts *in trans* to target the *HOXD* cluster for gene silencing (Rinn et al, 2007). Approximately 20 % of lncRNAs are associated with polycomb repressive complex 2 (PRC2) (Jing et al, 2010; Khalil et al, 2009), which is comprised of many subunits and functions to deposit histone H3K27 trimethylation (H3K27me3) and to suppress gene expression (Di Croce & Helin, 2013; Margueron & Reinberg, 2011; Simon & Kingston, 2013). Although some evidence indicates that lncRNAs might serve as scaffolds for PRC2 assembly and guide PRC2 to specific genomic targets, whether the interaction is specific and necessary in development or disease contexts is still unclear (Cifuentes-Rojas et al, 2014; da Rocha et al, 2014; Davidovich & Cech, 2015; Davidovich et al, 2015; Davidovich et al, 2013; Kaneko et al, 2014a; Kaneko et al, 2014b). Therefore, it is imperative to demonstrate that the specific interactions of lncRNAs with the PRC2 complex are functionally important and have specific regulatory targets to direct development or induce disease *in vivo*.

We used spinal motor neuron (MN) differentiation as a paradigm to assess these interactions. Although spinal cord development is one of the best characterized processes in the central nervous system (CNS) (Alaynick et al, 2011; Catela et al, 2016; Chen et al, 2011; Mazzoni et al, 2013a; Mazzoni et al, 2013b; Narendra et al, 2015; Philippidou & Dasen, 2013), how lncRNAs are involved in its transcription factor-driven gene regulatory networks is unclear (Briscoe & Small, 2015). MN differentiation into subtypes is mediated by the mutually exclusive expression of Hox transcription factors, which is programmed according to the body segment along the rostrocaudal (RC) axis. For example, segmental identity of MNs is defined by the mutually exclusive expression of Hox6, Hox9 and Hox10 (Dasen et al, 2003; Lacombe et al, 2013). In each segment, MNs are grouped into different columns according to their innervating targets. For instance, within the brachial Hox6^on^ segment, MNs are further grouped into axial muscle projecting MNs (Lhx3^on^, MMC) and forelimb-innervating MNs (Foxp1^on^, LMC). Finally, another set of mutually exclusive Hox proteins, such as Hox5 and Hox8 expression in the Foxp1^on^ LMC, further controls the rostral and caudal motor pool identity, which directs motor pools to either innervate proximal or distal muscles in the forelimb (Catela et al, 2016; Dasen et al, 2005b).

In the spinal cord, polycomb proteins control the exclusion of certain Hox protein expression at specific rostro-caudal (RC) positions and maintain this repression in differentiated cells. Depletion of the polycomb repressive complex 1 (PRC1) component Bmi1 at brachial level causes ectopic expression of Hoxc9 and subjects LMC neurons to a thoracic preganglionic column (PGC) fate. Conversely, elevation of Bmi1 represses Hoxc9 at thoracic level and subjects PGC neurons to an LMC fate (Golden & Dasen, 2012). These observations suggest that specific Hox repression may be maintained in MNs by distinct PRC1 activity levels, programmed along the RC axis. Recently, it was shown that during MN differentiation, *Hox* chromatin is demarcated into discrete domains controlled by opposing RC patterning signals (i.e., retinoic acid (RA), Wnt, and fibroblast growth factors (FGFs)) that trigger rapid and domain-wide clearance of H3K27me3 modifications deposited by PRC2 (Mazzoni et al, 2013b). More specifically, RA activates retinoic acid receptors (RARs) and binds to the *Hox1~5* chromatin domains, which is followed by synchronous domain-wide removal of H3K27me3 to acquire cervical spinal identity. At the tailbud, a gradient of Wnt and FGF signals induces expression of the Cdx2 transcription factor that binds and clears H3K27me3 from the *Hox1-Hox9* chromatin domains, thereby establishing brachial or thoracic segmental identity (Mazzoni et al, 2013b). Together, these findings indicate that epigenetic regulation of *Hox* clusters is critical to initiate and maintain patterns of *Hox* expression and that cross-repressive interactions of combinations of Hox proteins later consolidate the diversification of postmitotic MNs. However, the underlying mechanism that demarcates the histone modifiers at a molecular level is still unclear. Although many lncRNAs are known to regulate these histone modifiers, whether lncRNAs are directly involved in MN fate determination remains to be established.

We found that lncRNAs in the imprinted *Dlk1-Dio3* locus are highly enriched in postmitotic MNs. The *Dlk1-Dio3* locus contains three protein-coding genes (*Dlk1, Rtl1*, and *Dio3*) from the paternally inherited allele, and multiple lncRNAs and small ncRNAs are derived from the maternally inherited allele, including *Meg3, Rian* (containing 22 box C/D snoRNAs), as well as the largest miRNA mega-cluster in mammals (*anti-Rtl1*, which contains the *miR-127/miR-136* cluster of 7 miRNAs, and *Mirg* that contains the *miR-379/miR-410* cluster of 39 miRNAs). Interestingly, all of the ncRNAs are regulated by a common *cis*-element and epigenetic control, resulting in a presumable large polycistronic transcription unit (Das et al, 2015; Lin et al, 2003; Seitz et al, 2004). Although the *Dlk1-Dio3* locus is well known to play crucial roles in stem cells (Lin et al, 2007; Lin et al, 2003; Qian et al, 2016), we unexpectedly found that expressions of *Meg3* and other lncRNAs from the *Dlk1-Dio3* locus are also all enriched in postmitotic MNs. However, whether this locus functions during neural development had not been explored previously. Here, we show that lncRNAs in the imprinted *Dlk1-Dio3* locus shape postmitotic MNs by inhibiting progenitor and non-neural genes, and they also control MN subtype identity by regulating Hox expression. Our results provide strong evidence for the critical function of lncRNAs during MN development, emphasizing their physiological functions during embryonic development.

## RESULTS

### Identification of cell type-specific lncRNAs during MN differentiation

As epigenetic landscape remodelling and the cell fate transition during MN differentiation are well characterized (Chen et al, 2011; Li et al, 2017; Tung et al, 2015), we took advantage of an ESC differentiation approach that can recapitulate MN development to systematically identify cell type-specific lncRNAs during this differentiation process. Firstly, an ESC line harbouring the MN transgenic reporter *Hb9::GFP* was harnessed into MNs (Wichterle et al, 2002), and we then sequentially collected RA-induced nascent neural epithelia (Hoxa1^on^, NE at day 2), MN progenitors (Olig2^on^, pMN at day 4), and postmitotic MNs (Hb9::GFP^on^, postmitotic MNs at day 7) by fluorescence-activated cell sorting (FACS). Simultaneously, spinal interneurons (INs) derived from [smoothened agonist; SAG]^low^ conditions were collected and Hb9::GFP^off^ cells were sorted at day 7 as controls (Figure 1A). Next, we performed strand-specific RNA-seq across a library preserving non-polyadenylated transcripts while removing ribosomal RNAs, since many lncRNAs are non-polyadenylated (Yin et al, 2012; Zhang et al, 2012), and carried out *de novo* transcriptome assembly (Qian et al, 2016) to discover novel lncRNAs that might be specifically enriched during motor neuron development (detailed in the Supplemental Experimental Procedures and summarized in Figure 1B and Table S1). Several known markers for each cell type during MN differentiation were accurately recovered, corroborating the high quality and specificity of our RNA-seq data (Figure 1C). Our approach yielded 10,177 lncRNAs, 752 of which (7.39 %) were previously unidentified from the Ensemble mm10 database. We also found that 4,295 (77.78 %) of our identified lncRNAs overlapped with recently reported spinal MN-related lncRNAs, which were discovered by poly A^+^-enriched RNA-seq approaches (Amin et al, 2015; Narendra et al, 2015) (Figure S1A). Finally, we removed minimally expressed transcripts (TMM normalized read count < 10 in all samples), which left 602 expressed lncRNAs (Table S2). Based on stage-specific scores (see Supplemental Experimental Procedures), we identified 70 stage-signature lncRNAs during the ESC~MNs differentiation process (ESC, NE, pMN, MN, and IN in Figure 1D). Compared to protein-coding genes, both annotated lncRNAs and novel lncRNAs (newly identified in our *de novo* transcriptome assembly) had higher cell-type specificity (Figure 1E, Kolmogorov-Smirnov test, *P* = 1.41 × 10^−9^ and *P* = 3.53 × 10^−7^, respectively; see Supplemental Experimental Procedures), implying that lncRNAs might play specific roles in each cell type during ESC~MNs differentiation.

**Figure 1.**
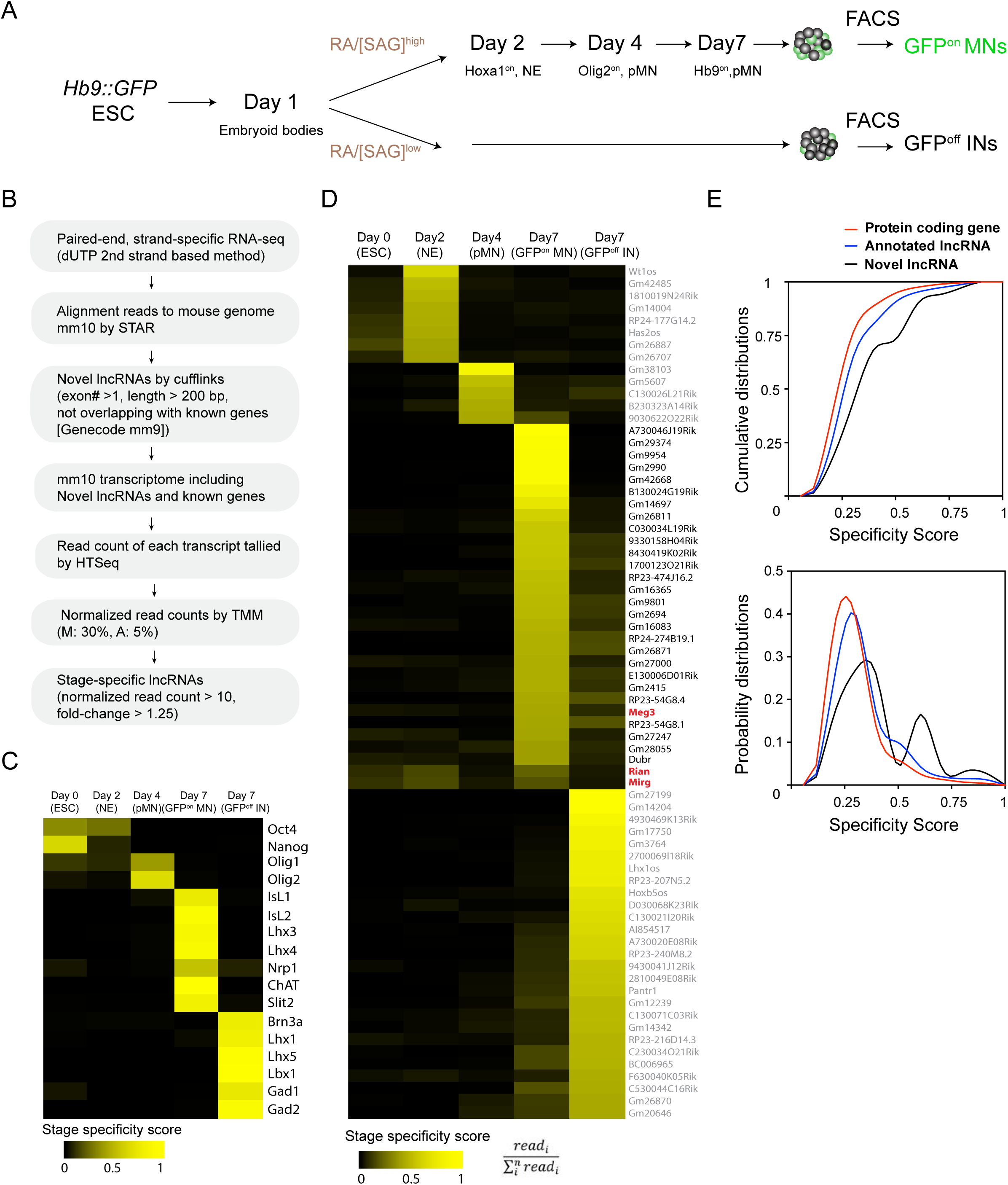
Identification of cell type-specific lncRNAs during motor neuron differentiation. (A) Schematic illustration of the differentiation process from *Hb9::GFP* ESCs to spinal MNs. RA: retinoic acid. SAG: Smoothened agonist. ESC: embryonic stem cell. NE: neural epithelium. pMN: motor neuron progenitor. MN: motor neuron. IN: interneuron. (B) Workflow of the RNA-seq analysis to identify cell type-enriched lncRNAs. (C and D) Heatmaps presenting the abundances of known cell type-specific transcription factors (C) and the abundances of lncRNA signatures (D) across each stage from ESCs to postmitotic MNs and INs (color indicates specificity scores). (E) Cumulative distributions (above) and probability distributions (below) of the stage specificity score of different categories of genes (protein coding genes [red], annotated lncRNAs [blue] and novel lncRNAs [black]), representing a measure of differential expression for each transcript across the cell types. The distribution reveals that annotated lncRNAs and novel lncRNAs manifest significantly higher specificity (according to Kolmogorov-Smirnov tests) than protein coding genes.

### *Dlk1-Dio3* locus-derived lncRNAs are enriched in the nuclei of postmitotic MNs

To identify developmentally up-regulated lncRNAs, we compared day 4 pMNs vs. day 7 postmitotic MNs (Figure 2A). Furthermore, to differentiate cell type-specific lncRNAs, we performed a pairwise comparison of day 7 postmitotic MNs against day 7 INs (Figure 2B). We identified 117 lncRNA candidates from our analysis as being postmitotic MN-enriched. We further selected several MN-lncRNAs that manifested high normalized reads from RNA-seq data and verified their MN-specific expression by qPCR (Figure S1B). Interestingly, the lncRNAs *Meg3, Rian*, and *Mirg*, which are transcribed from the imprinted *Dlk1-Dio3* locus on mouse chromosome 12qF, all manifested strong enrichment in postmitotic MNs (Figure S2A). Since these lncRNAs are conserved amongst placental mammals (Ogata & Kagami, 2016), we chose to characterize their functions in greater detail.

**Figure 2.**
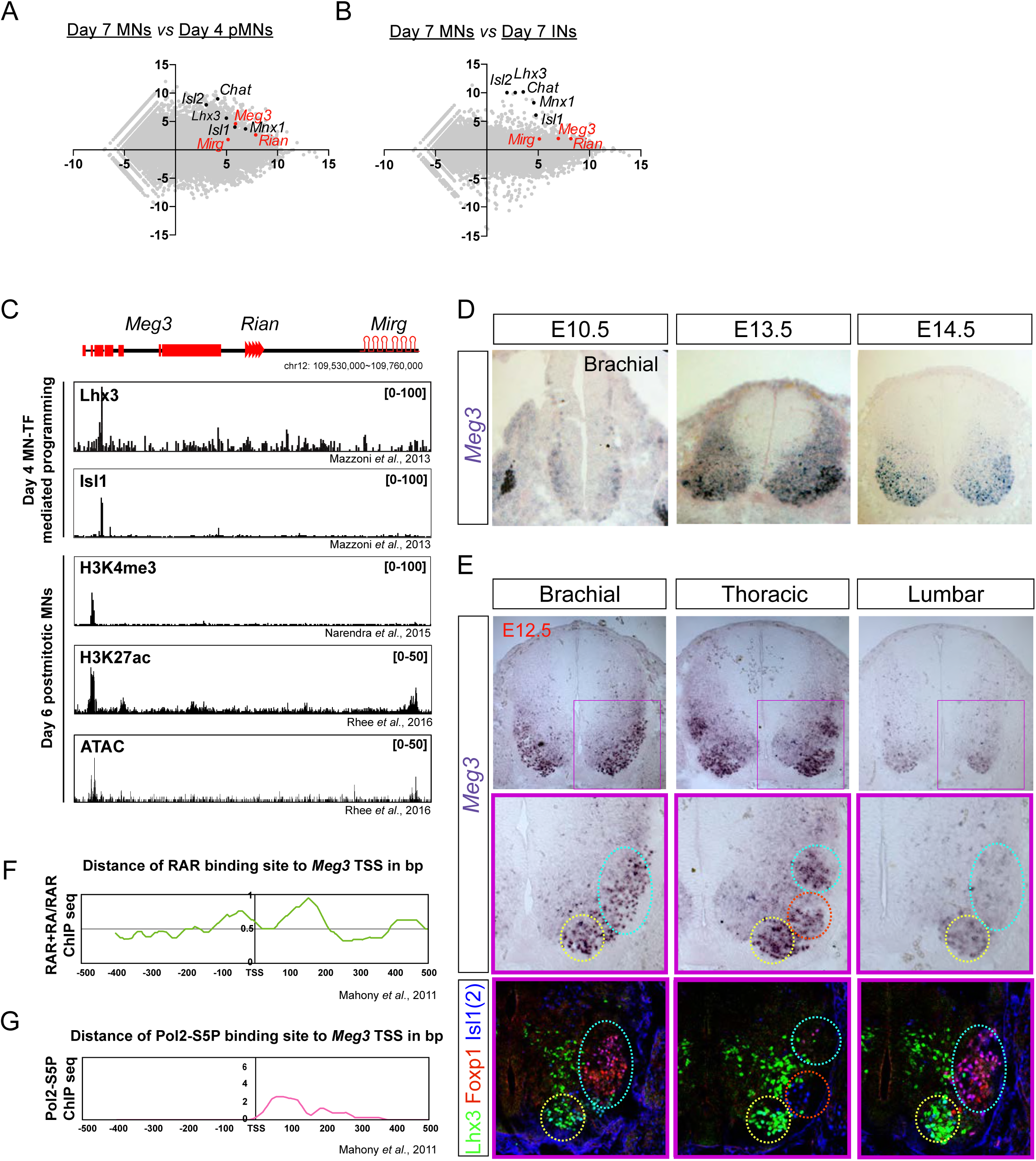
*Dlk1-Dio3* locus-derived lncRNAs are enriched in postmitotic motor neurons. (A and B) MA plots demonstrate that *Meg3*, *Rian*, and *Mirg* are postmitotic (day 7 MNs vs. day 4 pMNs; A) MN signature (day 7 MNs vs. day 7 Ins; B) lncRNAs. X-axis: mean abundance; Y-axis: log_2_ fold-change. (C) Illustration of the imprinted *Dlk1-Dio3* locus. The lncRNAs of the maternally-inherited allele (in red) are on mouse chromosome 12. The miRNA and C/D snoRNA genes are shown by hairpin loops and triangles, respectively. Comparison of ChIP-seq for MN transcription factors (Lhx3 and Isl1), H3K4me3, and H3K27ac, together with ATAC-seq of the *Dlk1-Dio3* locus. (D) *In situ* hybridization shows that *Meg3* is gradually enriched and restricted in postmitotic MNs in the developing spinal cord. (E) *In situ* hybridization together with adjacent sections of immunostaining on E12.5 segmental spinal cords demonstrate that *Meg3* is enriched at brachial and thoracic MNs (Isl1/2^on^), but no preference for columnar MN subtypes was revealed for Foxp1 or Lhx3. (F and G) RAR binding and recruitment of the elongating form of Pol2 to the *Meg3* transcription start site (TSS) occur within 8 hours of retinoic acid (RA) exposure.

To investigate why *Meg3-Rian-Mirg* are highly enriched in postmitotic MNs, we examined the binding landscape of MN-specific transcription factors (i.e., Lhx3 and Isl1), chromatin accessibility (ATAC-seq), and histone modification (H3K4me3 and H3K27ac) across the *Meg3-Rian-Mirg* locus from previously published studies (Figure 2C) (Mazzoni et al, 2013a; Narendra et al, 2015; Rhee et al, 2016). Within this locus, we uncovered an MN-specific active chromatin region that possesses enhancer/promoter characteristics with direct MN-specific transcription factor binding (Figure 2C). Furthermore, overexpression of MN-TFs in a maternally-inherited intergenic differentially methylated region (*IG-DMR*) deletion (*IG-DMR^matΔ^*) ESC line, which leads to simultaneous silencing of all maternally-expressed lncRNAs in the *Meg3-Rian-Mirg* locus but leaves the MN-TF binding site intact (Figure S2A) (Lin et al, 2007; Lin et al, 2003), can robustly induce *Meg3* expression (Figure S2B). Therefore, we suggest that MN-TFs bind and directly activate *Meg3-Rian-Mirg* during differentiation.

We further performed *Meg3 in situ* hybridization and immunostaining of the adjacent sections along the RC axis from E10.5~12.5. We found that *Meg3* expression: 1) is enriched in the mantle zone of the developing spinal cord during development and is gradually enriched in postmitotic MNs (Isl1/2^on^ cells) after E12.5; 2) has no preference for columnar MN subtypes, as revealed by Foxp1 (LMC-MNs) and Lhx3 (MMC-MNs) immunostaining; and 3) exhibits rostral high (brachial and thoracic) and caudal low (lumbar) asymmetry after E12.5 (Figures 2D and 2E).

Why does *Meg3* exhibit strong enrichment in the brachial spinal cord? Given that previous reports indicate that rostral *Hox* genes enriched in the brachial spinal cord are mediated by an RA gradient (Mazzoni et al, 2013b; Novitch et al, 2003), we hypothesized that *Meg3* might also be induced by RA. To examine this possibility, we checked if there is any RA-driven binding to RAR sites near the *Meg3* promoter (Figure 2F) (Mahony et al, 2011). Interestingly, we found that RA treatment results in novel binding of RAR directly to the *Meg3* promoter, as well as subsequent recruitment of the basal transcription complex (Pol2 S5P in Figure 2G). Moreover, we observed that *Meg3* is induced after the addition of RA in *IG-DMR^matΔ^* ESCs after 8 hours (Figure S2C), indicating that RA/RAR activation triggers the strong *Meg3* expression in rostral brachial MNs.

Finally, to characterize the abundance and subcellular localization patterns of *Meg3* at a cellular level, we designed a set of single molecule RNA FISH probes specific to *Meg3* and examined their expression in ESC~MNs. We observed a speckled pattern of *Meg3* expression enriched in the nucleus, suggesting it has a potential function in gene regulation (Figure S2D). Furthermore, qPCR of subcellular-fractionated RNAs from ESC~MNs validated that *Meg3* is not only enriched in the nucleus, but that it is also chromatin-associated (Figure S2E; *Gapdh* as cytoplasmic marker, *U1* as nuclear marker, and *Kcnq1ot1* as a chromatin-associated RNA control). Together, these findings suggest that lncRNAs in the *Dlk1-Dio3* locus are postmitotic MN-enriched, and that they are directly activated by MN-TFs and RA/RAR. At a cellular level, *Meg3* is highly enriched in MN nuclei and is chromatin-associated, indicating a potential function in chromatin regulation.

### *Meg3* facilitates interaction of the PRC2 complex with Jarid2 in MNs

Several previous reports have revealed that interactions between Jarid2 and ncRNAs regulate PRC2 recruitment to chromatin, including lncRNAs in the *Dlk1-Dio3* locus of ESCs (da Rocha et al, 2014; Kaneko et al, 2014a; Kaneko et al, 2014b). The roles of PRC2/Jarid2 in postmitotic cells are less clear. Unlike the PRC2 complex, Jarid2 is known to have diverse cell type-specific functions (Landeira & Fisher, 2011). Surprisingly, we found that expression of *Jarid2* was reactivated in postmitotic MNs, pointing to specific regulation in this cell type (Figure S3A) (Takeuchi et al, 1995). This prompted us to examine if lncRNAs in the *Dlk1-Dio3* locus bind to the PRC2 complex and maintain postmitotic MN fate by controlling the H3K27me3 landscape. To test this hypothesis, we first demonstrated that immunoprecipitation (IP) of endogenous PRC2 complex components (i.e., Ezh2 and Suz12), as well as the PRC2 cofactor Jarid2, from ESC~MNs specifically retrieves *Meg3, Rian* and *Mirg* RNA, whereas the nuclear ncRNA *U1* and the lncRNA *Malat1* were not captured by Ezh2, Jarid2, or Suz12 (Figure 3A). Given that *Rian* and *Mirg* are further processed to snoRNAs and miRNAs (Lin et al, 2003) and that *Meg3* is known to regulate pluripotency (Stadtfeld et al, 2010), imprinting (Das et al, 2015), and PRC2 function (Zhao et al, 2010), we focused on biochemical characterization of *Meg3*. Several *Meg3* isoforms have previously been documented, so we scrutinized across the entire *Meg3* locus (~31 kb) and found that two isoforms, *Meg3^v1^* and *Meg3^v5^* (Ensemble mm10 database), are predominantly expressed in Hb9::GFP^on^ MNs (Figures S3B and S3C). This result was confirmed by qPCR, which indicated that expression of *Meg3^v5^* is more significantly enriched in Hb9::GFP^on^ MNs compared to the other common variants that have previously been studied in ESC~MNs (Figure S3D) (Kaneko et al, 2014a; Zhou et al, 2007). Interestingly, *Meg3^v1^* and *Meg3^v5^* have mutually exclusive exon sequences (Figure S3E), raising the possibility that the two isoforms might exert different functions. However, both purified biotinylated *Meg3^v1^* and *Meg3^v5^* RNA retrieved Ezh2 from cell nuclear extracts of ESC~MNs (Figure 3B; *GFP* RNA was used as a negative control). These results suggest that these *Meg3* isoforms directly interact with PRC2/Jarid2 complexes and might facilitate association of the PRC2 complex with Jarid2.

**Figure 3.**
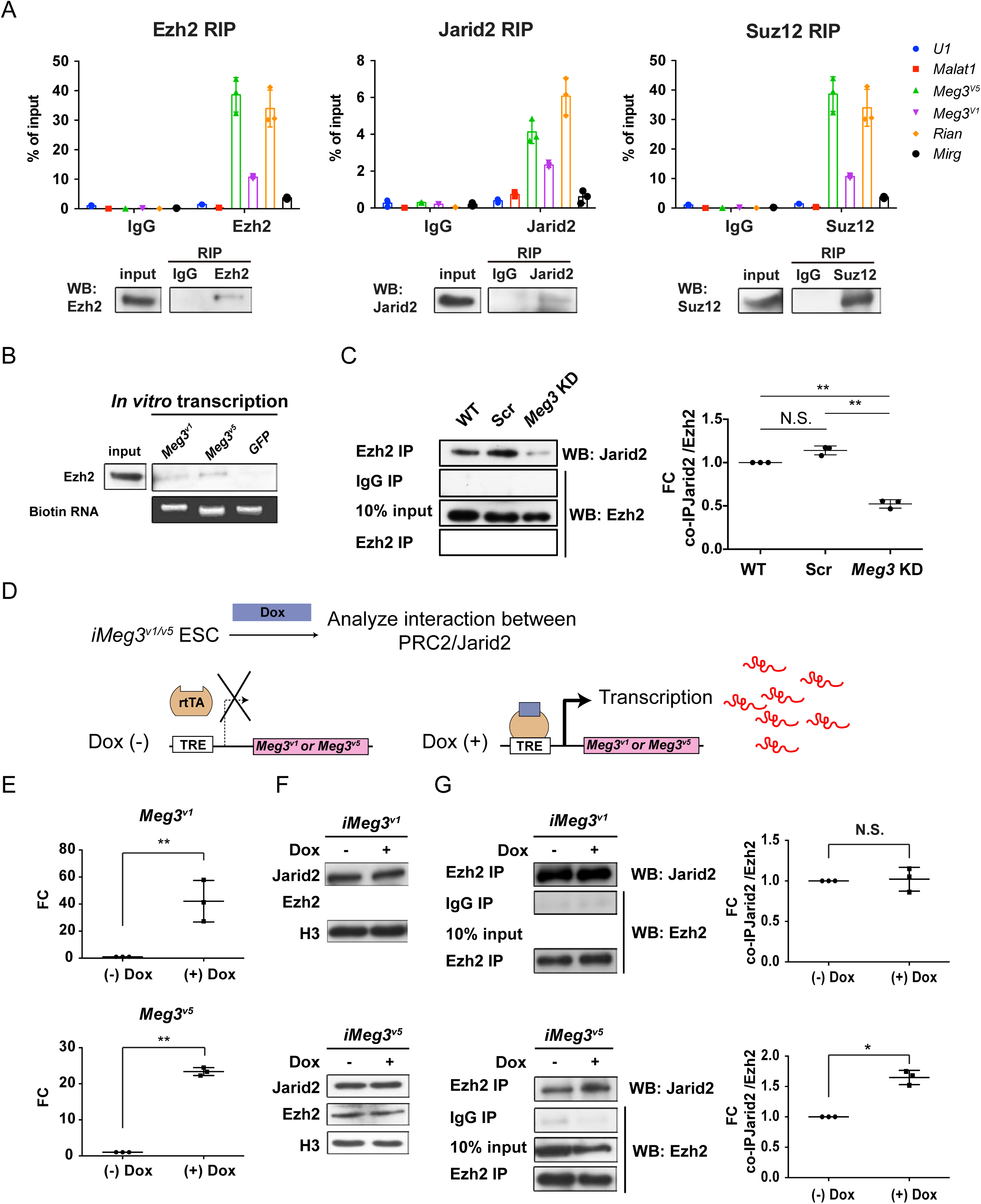
*Meg3* facilitates the non-stoichiometric interaction of the PRC2 complex and Jarid2. (A) Ezh2, Jarid2, and Suz12 immunoprecipitation specifically retrieves *Meg3* RNA isoforms (v1 and v5). *U1* RNA and *Malat1* lncRNA are negative controls. 10 % input was used to normalize the retrieval efficiency (error bars represent SD, n=3 independent experiments). Immunoblotting reflects the recovery of Ezh2, Jarid2 and Suz12 proteins using the corresponding antibodies. (B) *In vitro*-transcribed (IVT), biotinylated *Meg3* RNA isoforms retrieved Ezh2. (C) Ezh2 interacts with Jarid2 in ESC~MNs, but knockdown of *Meg3* impairs this interaction. The abundance of Jarid2 is shown on the right (N.S.: not significant; error bars represent SD, n=3 independent experiments; ** p-value < 0.01 by Student’s *t*-test). (D and E) The design of inducible “Tet-On” ESC lines expressing *Meg3^v1^* or *Meg3^5^* under the doxycycline (Dox)-regulated promoter. In the presence of Dox, the reverse tetracycline-controlled transactivator (rtTA) is recruited to the TRE (tetracycline response element), thereby initiating robust transcription of *Meg3^v1^* or *Meg3^v5^*, respectively. (F and G) Overexpression of *Meg3^v1^* or *Meg3^v5^* does not alter the protein levels of Ezh2 or Jarid2. *Meg3^v5^* but not *Meg3^v1^* stimulates more Ezh2 and Jarid2 interaction. The abundances of Jarid2 are shown on the right (FC: fold-change; N.S.: not significant; error bars represent SD, n=3 independent experiments; * p-value < 0.05, ** p-value < 0.01 by Student’s *t*-test).

As the PRC2 complex and Jarid2 are known to interact in a non-stoichiometric manner (Pasini et al, 2010; Peng et al, 2009), we further examined if *Meg3* facilitates the interaction between PRC2 complex and Jarid2. To test this possibility, we performed IP with Ezh2 (a core component of PRC2) to retrieve Jarid2 from ESCs (Figure 3C). We first verified that *Meg3* knockdown (KD) did not affect the protein abundance of Ezh2/Jarid2, but we did observe that it undermined the interaction between Ezh2 and Jarid2, suggesting that *Meg3* facilitates this interaction (Figure 3C and Figure S3F). We then investigated if the two *Meg3* isoforms have differing abilities to facilitate Ezh2/Jarid2 binding by generating two locus-defined Tet-ON-inducible *Meg3* ESCs (Figure 3D, *iMeg3^v1^* and *iMeg3^v5^*). Upon doxycycline induction, both *Meg3* isoforms were induced ~20–50 fold, yet the abundance of Ezh2/Jarid2 remained unaffected (Figures 3E and 3F). *Meg3^v5^* overexpression in ESCs significantly increased the binding of Ezh2 and Jarid2, whereas *Meg3^v1^* overexpression had a minimal effect (Figure 3G), indicating that *Meg3^v5^* is a strong facilitator of the binding of the PRC2 complex and Jarid2. Accordingly, we suggest that *Meg3*, and particularly the *Meg3^v5^* isoform, facilitates the binding of the PRC2 complex and Jarid2 in postmitotic MNs.

#### *Dlk1-Dio3* locus-derived lncRNAs maintain the epigenetic landscape in postmitotic MNs

To test if the binding of Ezh2/Jarid2 by the lncRNAs in the *Dlk1-Dio3* locus is important to maintain the epigenetic landscape in postmitotic MNs, we systematically analyzed genome-wide H3K27me3 profiles of control and *IG-DMR^matΔ^* ESC~MNs by ChIP-seq (chromatin immunoprecipitation-sequencing) (Figure 4A). To overcome the complication of concomitant up-regulation of paternal coding genes in *IG-DMR^matΔ^* ESCs (Lin et al, 2007; Lin et al, 2003), we further established two retrovirus-based short hairpin RNAs (shRNAs) targeting *Meg3* and used a knockdown approach to prevent impairment of DMR sites. Both shRNAs reduced the expression of *Meg3* by an average of ~90 % compared to endogenous levels in ESC~MNs (Figure S4A). As negative controls, we performed independent infections with retroviruses containing scrambled shRNA with no obvious cellular target RNA. We selected two stable *Meg3* knockdown (KD) ESCs (referred to as H6 and K4 hereafter) that had the best ESC morphology for further experiments. Verification by qPCR indicated that the expressions of two other lncRNAs, *Rian* and *Mirg*, from the maternal allele of the *Dlk1-Dio3* imprinted locus were all significantly down-regulated in *Meg3* KD MNs, whereas paternal genes were unaffected (Figure S4A). This finding is consistent with a previous report indicating that *Meg3-Rian-Mirg* probably represents a single continuous transcriptional unit (Das et al, 2015).

**Figure 4.**
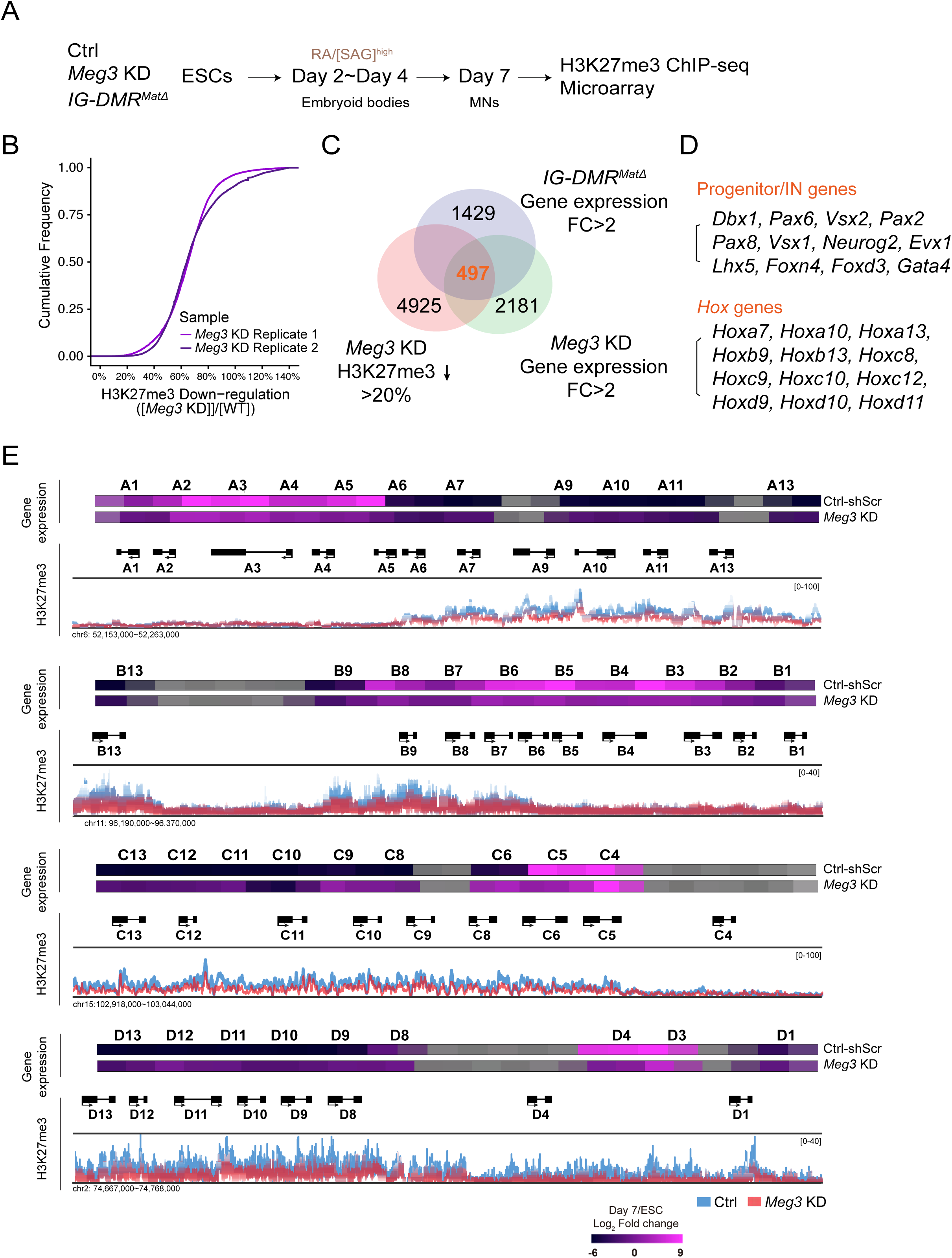
Loss of *Dlk1-Dio3* locus-derived lncRNAs in MNs leads to dysregulation of neural progenitor and *Hox* genes. (A) Genome-wide profiling of H3K27me3 by ChIP-seq and gene expression by Agilent microarray in control, *IG-DMR^matΔ^*, and *Meg3* KD ESC~MNs. Schematic illustration of the MN differentiation process from control scramble-shRNA-targeted ESCs (Ctrl-shScr) and two independent *Meg3* shRNA-mediated KD ESCs (H6 and K4). (B) Cumulative plot reflects a global trend of down-regulation of H3K27me3 level in two independent *Meg3* KD derived MNs. (C) Venn diagram shows the number of genes that are up-regulated in both *IG-DMR^matΔ^* and *Meg3* KD MNs while also displaying down-regulation of the H3K27me3 epigenetic landscape. (D) Loss of *Meg3* imprinted lncRNAs is related to the up-regulation of progenitor and spinal interneuron genes, as well as *Hox* regulations. (E) Heatmaps illustrating the expression profiles of ESCs and ESC~MNs in control scrambled and *Meg3* KD lines. The color indicates the log_2_ fold-change in signal intensity between ESCs and ESC~MNs. Genes in light grey are not represented in the microarrays. Loss of *Meg3* imprinted lncRNAs results in ectopic expression of a majority of caudal *Hox* genes (*Hox8~13*), with concomitant down-regulation of the H3K27me3 levels.

We then performed H3K27me3 ChIP-seq of control and *Meg3* KD MNs. We observed a trend of global down-regulation of H3K27me3 in both independent experiments of *Meg3* KD MNs, most likely a reflection of compromised Ezh2/Jarid2 interaction (Figure 4B and Figure S4B). Since the response to PRC2 activity change in a given cell type might be context-dependent (Davidovich et al, 2013), we sought to identify the most prominent genes in MNs regulated by *Dlk1-Dio3* locus-derived lncRNAs based on the loss of H3K27me3. To achieve this, we profiled gene transcriptomes of control, *IG-DMR^matΔ^*, and *Meg3* KD MNs. Next, we compared the co-upregulated genes between *IG-DMR^matΔ^* and *Meg3* KD MNs, together with H3K27me3 landscape upon the loss of ncRNAs in the *Dlk1-Dio3* locus (Figure 4C and FigureS4C). This approach revealed 497 genes in MNs that displayed down-regulation of the H3K27me3 epigenetic landscape and concomitant up-regulation of gene expression upon loss of the *Meg3* imprinted lncRNAs (Figure 4C). Gene ontology (GO) analysis of these genes revealed significant enrichment for rostrocaudal patterning and progenitor genes, and strikingly so for homeodomain *Hox* genes (Figures 4D and Figure S4D; false discovery rate (FDR) q-value ≤ 0.05). We observed that a majority of caudal *Hox* genes (*Hox8~13*) were up-regulated with a concomitant down-regulation of the H3K27me3 epigenetic landscape (Figure 4E). We corroborated this finding by generating a third *Meg3* KD ESC line (I6) and confirming that all *Meg3* KD ESC~MNs exhibited imbalanced expression of 3’ and 5’ *Hox* genes across entire *Hox* clusters (Figure S4E). Thus, for both *Meg3* KD and *IG-DMR^matΔ^* MNs, dysregulation of progenitor and *Hox* gene expression is the most prominent phenotype, likely due to loss of the robustness of the epigenetic landscape of postmitotic MNs.

#### *IG-DMR^matΔ^* embryos manifest dysregulation of progenitor and *Hox* genes in postmitotic MNs

To corroborate the observed phenotype of *IG-DMR^matΔ^* ESC~MNs, we further scrutinized the MN phenotype in *IG-DMR^matΔ^* embryos. Consistent with previous studies, *IG-DMR^matΔ^* embryos died soon after E16 (Lin et al, 2007), so we analyzed MN phenotypes from E10.5~E13.5 in this study. Since we had observed that the progenitor gene *Dbx1* is one of the most up-regulated progenitor genes in *IG-DMR^matΔ^* ESC~MNs (Figure S4C), we first checked if *Dbx1* is affected by crossing *IG-DMR^matΔ^* to the *Dbx1^LacZ/+^* reporter mice (Bielle et al, 2005). Compared to the control *Dbx1^LacZ/+^* littermates, we observed a significant increase in the number of Dbx1^on^ cells along the entire rostrocaudal axis of the ventral spinal cord in *IG-DMR^matΔ^; Dbx1^LacZ/+^* embryos (Figures 5A and 5C, only the cervical segment is shown). However, Hb9^on^ and Isl1(2)^on^ MNs were comparable between control and *IG-DMR^matΔ^* embryos at E10.5 (Figures 5B and 5C). This finding indicates that although progenitor genes are aberrantly up-regulated in the ventral spinal cord, production of MNs remains relatively unaffected.

**Figure 5.**
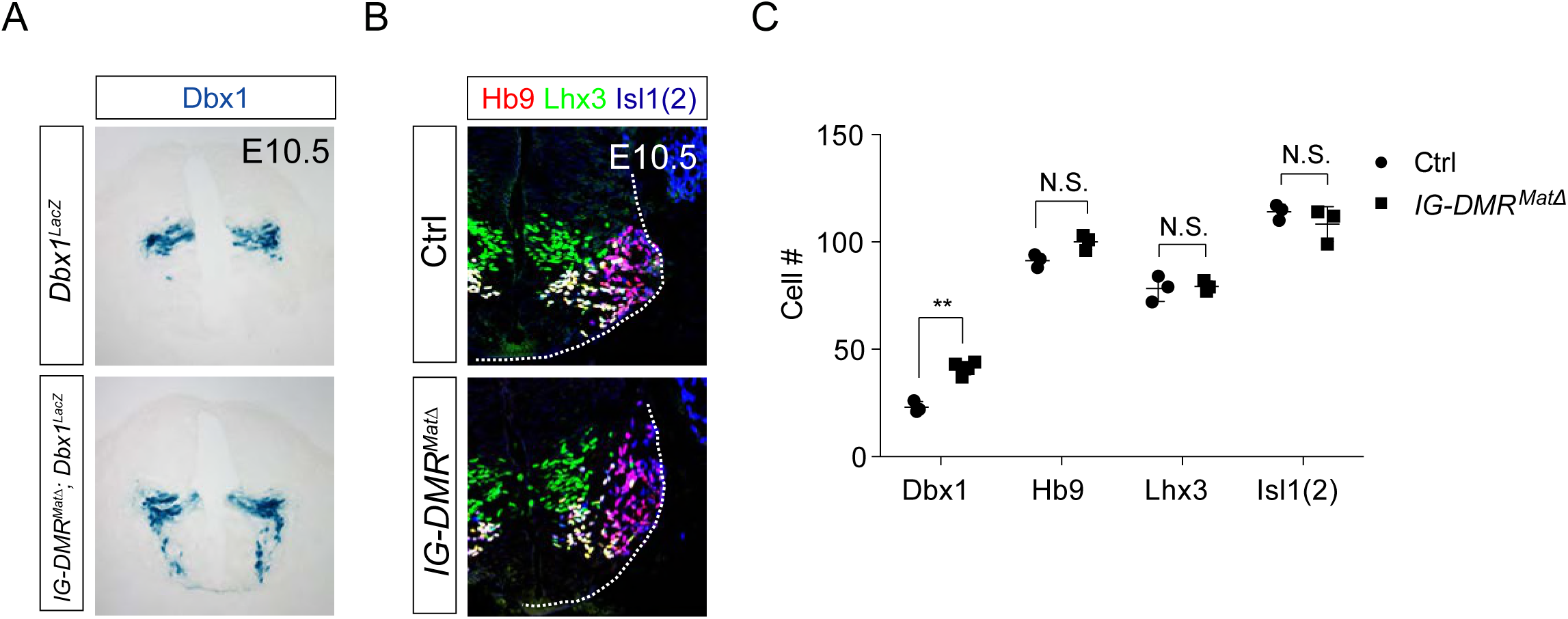
*IG-DMR^matΔ^* mice ectopically turn on Dbx1 expression in the ventral spinal cord. (A) *IG-DMR^matΔ^* embryos display ectopic Dbx1^on^ cells in the ventral spinal cord, as revealed by X-gal staining. (B) Specification of generic MNs (Hb9^on^, Lhx3^on^, and Isl1(2)^on^) is not affected in spinal cords of *IG-DMR^matΔ^* mice at E10.5. Quantification of postmitotic MNs (number of positive cells per 15 µm brachial ventral-half sections) in wild type control and *IG-DMR^matΔ^* embryos (error bars represent SD, n=3 embryos at E10.5; N.S.: not significant; * p-value < 0.05, ** p-value < 0.01 by Student’s *t*-test).

Next, we checked if the expression of Hox proteins is affected in *IG-DMR^matΔ^* embryos. We first assessed how loss of *Dlk1-Dio3* locus-derived lncRNAs affected the specification of segmental MNs, marked by brachial (Hoxc6), thoracic (Hoxc9), and lumbar (Hoxd10) Hox levels. We observed comparable numbers of cells expressing respective Hox proteins at each segmental level between control and mutant embryos (Figures S5A and S5B). Columnar identities of axial (Lhx3^on^) and limb-innervating MNs (Foxp1^on^) were also largely unaffected in the *IG-DMR^matΔ^* embryos (Figures S5C and S5D).

To further examine MN subtype diversification within the limb-innervating MNs, we checked the Hox proteins involved in pool specification (Catela et al, 2016; Dasen et al, 2005a). Whereas reciprocal expression of Hoxa5 and Hoxc8 was maintained along the rostrocaudal axis in the Hox6^on^ LMC MNs of control embryos, Hoxc8 was expanded rostrally into Hoxa5^on^ territory in *IG-DMR^matΔ^* embryos, along with a significant increase of the Hoxc8-mediated downstream motor pool genes, Pea3 and Scip (n=5 embryos in Figures 6A and 6C for rostral brachial segments and Figures 6B and 6D for caudal brachial segments; quantification in Figures 6E and 6F). The reduction of Hoxa5 was not attributable to apoptosis, as cCasp3^on^ cells were comparable in both control and *IG-DMR^matΔ^* mutant embryos (data not shown).

**Figure 6.**
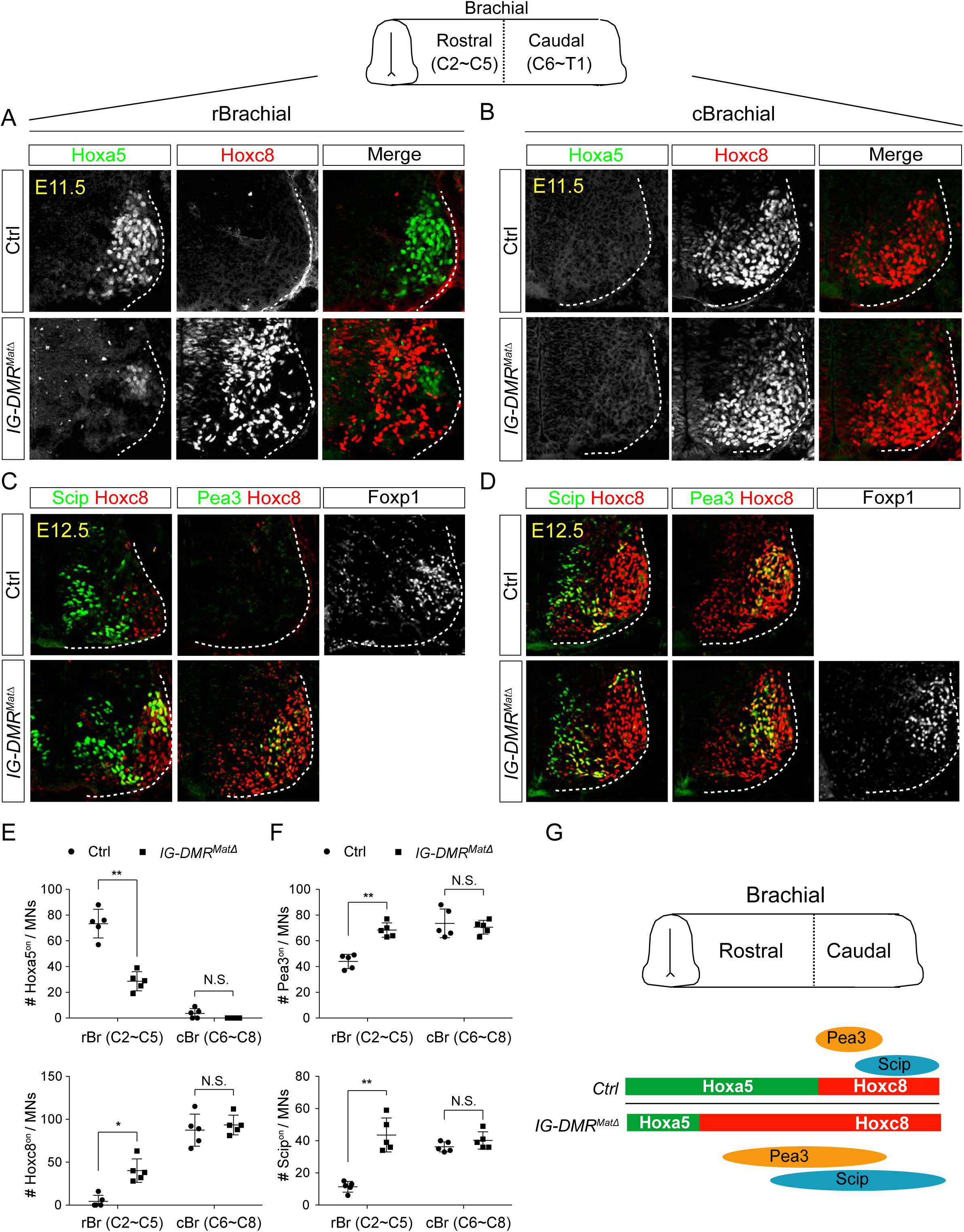
*IG-DMR^matΔ^* mice manifest switched MN subtype identity. (A~D) Ectopic expansion of Hoxc8 and downstream Pea3^on^ and Scip^on^ MN pools in the rostral brachial segment, with a concomitant decrease of Hoxa5^on^ MNs for E11.5~E12.5 *IG-DMR^matΔ^* embryonic spinal cord sections. Expression of Hoxc8 in the caudal brachial region remains unaffected (B and D). (E and F) Quantification of the numbers and distributions of Hoxa5^on^, Hoxc8^on^, Pea3^on^, and Scip^on^ MNs in the control and *IG-DMR^matΔ^* mice from serial sections along the rostrocaudal axis (N.S.: not significant; error bars represent SD, n=5; * p-value < 0.05, ** p-value < 0.01 by Student’s *t*-test). (G) Summary of the motor neuron phenotype in *IG-DMR^matΔ^* embryos.

### Peripheral innervation defects in *IG-DMR^matΔ^* mutants

To further examine the impact of switching the MN pool subtype identity of LMC-MNs, we assessed the potential trajectory and target selectivity of motor axons in wild type control and *IG-DMR^matΔ^* embryos. We bred *IG-DMR^matΔ^* mutants to a transgenic line of *Hb9::GFP* mice in which all motor axons are labeled with GFP and then analyzed the overall pattern of limb innervation. First, the images of motor nerves from light sheet microscopy were converted into panoramic 3D images (Figure 7A upper panel; Supplementary Movies 1 and 2). The overall trajectory of each motor nerve was reconstructed by Imaris (lower panel in Figure 7A) and this conversion enabled semi-automatic calculation of the number of motor nerve terminals in each skeletal muscle, as well as comparison of the extent of motor axon arborization between skeletal muscles (see Supplementary Experimental Procedures for details). Under higher magnification with better resolution, we observed the terminal arbors of suprascapular (Ss) nerves of scapulohumeralis posterior muscles were significantly eroded and reduced (Figure 7B), consistent with the caudalized switch from Hoxa5 to Hoxc8 expression within LMC neurons. Concomitantly, increased arborization complexity of distal muscle-innervating nerves, including axillary (Ax) and posterior brachial cutaneous (PBC) nerves were manifested in the *IG-DMR^matΔ^* embryos (Figures 7C and 7D, quantification in 7E, n = 6, P <0.01, Mann-Whitney U test). Thus, the *IG-DMR^matΔ^* mutants displayed deficiencies in peripheral innervation of MNs, which might be a consequence of dysregulation of Hox proteins and/or other axon arborization genes in the absence of lncRNAs from the *Dlk1-Dio3* locus.

**Figure 7.**
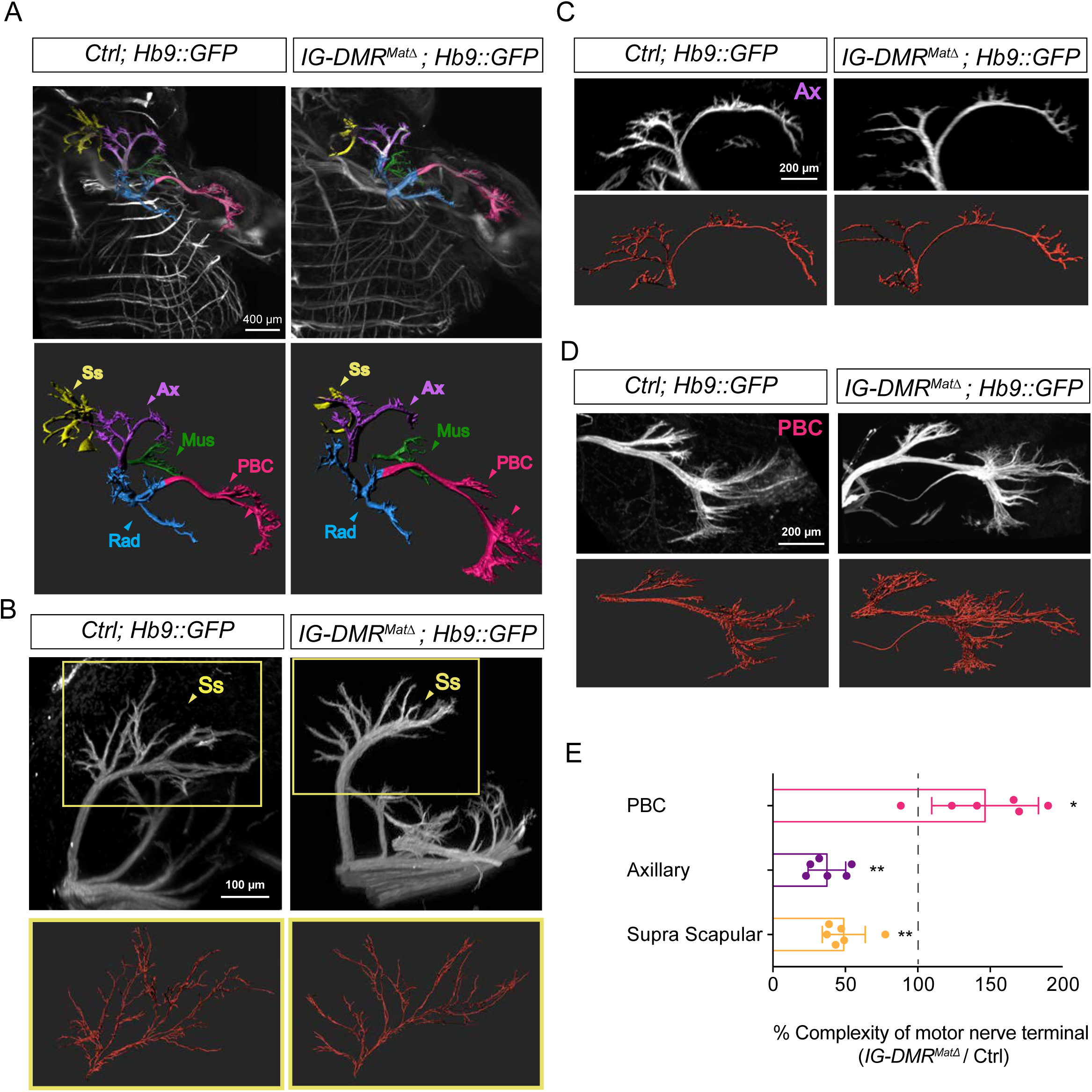
Motor axon innervation defects in *IG-DMR^matΔ^* mutants. (A) Panoramic views from different angles of control and *IG-DMR^matΔ^*; *Hb9::GFP* embryos at E13.5 using light sheet microscopy (Upper panel). Details of each viewing angle are illustrated in Supplementary Movies 1 and 2. Reconstruction of motor nerve positions by Imaris (see supplementary methods for details) is illustrated in the lower panel. Suprascapular nerve (Ss, yellow); axillary nerve (Ax, purple); musculocutaneous nerve (Mus, green); radial nerve (Rad, blue); posterior brachial cutaneous nerve (PBC, pink). (B~D) Higher magnification of MN innervations in the forelimbs of E13.5 control and *IG-DMR^matΔ^*; *Hb9::GFP* mice. Mutant mice display defects in Ss and Ax axonal branching, concomitant with more PBC axonal branching. Semi-automatic highlighting of the axonal branching and nerve trajectories is used and quantified by Imaris. (E) Quantification of the axonal branching and nerve trajectories for E13.5 control and *IG-DMR^matΔ^*; *Hb9::GFP* mice by Imaris (see Supplementary Experimental Procedures for details) (n = 6, p < 0.01, Mann–Whitney U test).

## DISCUSSION

Although mammalian genomes encode tens of thousands of lncRNAs, only less than a hundred have been shown to play critical roles in gene regulation *in vitro*. Consequently, the *in vivo* functions of the vast majority of lncRNAs remain to be vigorously tested. Strikingly, 40 s of lncRNAs are expressed specifically in the central nervous system (CNS), which makes it one of the best systems for uncovering the physiological functions of lncRNAs (Briggs et al, 2015; Ng et al, 2013). In this study, we identified a series of novel and/or uncharacterized lncRNAs that exhibit precisely regulated temporal and spatial expression patterns during MN development. Here, we focused on characterizing lncRNAs located in the imprinted *Dlk1-Dio3* locus for three reasons: 1) this locus is conserved between human and mouse (Lin et al, 2007; Lin et al, 2003); 2) several studies have highlighted that *Meg3* might function as a tumor suppressor (Zhang et al, 2010; Zhou et al, 2007); and 3) a previous report indicates that the paternally-expressed coding gene *Dlk1* has an unexpected function in determining MN subtype diversification (Muller et al, 2014), which prompted us to examine whether the MN-enriched lncRNAs in the same locus also participate in MN cell fate determination.

### Functional perspectives of lncRNAs in MNs

Although lncRNAs derived from the *Dlk1-Dio3* locus are highly expressed in the CNS, their functions during neural development are largely unknown (Wang et al, 2012; Zhang et al, 2003; Zhou et al, 2012). Upon knockdown of *Meg3* (a *Dlk1-Dio3* locus-derived lncRNA), we uncovered that: 1) many adjacent progenitor genes were significantly up-regulated; and 2) the rostral *Hox* genes were significantly down-regulated, with a concomitant increased expression of caudal *Hox* genes in ESC~MNs. This phenotype was recapitulated in *IG-DMR^matΔ^* embryos, in which Hoxc8 expression is expanded in Hoxa5^on^ MNs. Several reports have identified that certain lncRNAs can shape the *Hox* epigenetic landscape by *cis* and *trans* modulation (Dasen, 2013; Rinn et al, 2007; Wang et al, 2011). In addition, we recently uncovered that a novel *trans* Hox-miRNA circuit can filter *Hox* transcription noise and control the timing of protein expression to confer robust individual MN identity (Li et al, 2017). Here, we have now added the *Meg3* imprinted lncRNA to that list as a novel trans-acting lncRNA that maintains the *Hox* epigenetic landscape, most likely by recruiting Jarid2 to the PRC2 complex.

### PRC1/2 with lncRNA: a fail-safe mechanism to guard MN epigenetic landscape

Why do MNs deploy an lncRNA-mediated strategy to maintain postmitotic cell fate by inhibiting progenitor genes and regulating Hox boundaries? The dynamic role of lncRNAs in modulating PRC2 function is well documented; ranging from recruitment, complex loading and activity control to gene targeting (Davidovich & Cech, 2015; Kretz & Meister, 2014). A recent study of Drosophila *Hox* genes revealed that epigenetic H3K27me3 chromatin modification functions as a legitimate carrier of epigenetic memory (Coleman & Struhl, 2017), providing compelling evidence for a physiologically significant role of chromatin modification in epigenetic inheritance. Nonetheless, the epigenetic memory carried by H3K27me3 in a postmitotic cell may still be overridden by H3K27me3-opposing demethylases (Coleman & Struhl, 2017). Our RNA-seq data revealed that two prominent H3K27me3 demethylases, *Kdm6a (Utx)* and *Kdm6b (Jmjd3)*, are reactivated in postmitotic MNs (data not shown). Although the function of H3K27me3 demethylase reactivation in postmitotic MNs remains unknown, these findings raise the possibility that the epigenetic memory of the H3K27me3 landscape in postmitotic MNs might still need to be “actively” maintained to counterbalance H3K27me3 demethylase activity. Enrichment of postmitotic MN lncRNAs might therefore bridge and scaffold the PRC2/Jarid2 interaction and activity to maintain MN epigenetic memory by repressing progenitor genes and also carve MN subtype identity by repressing caudal *Hox* genes (Figure S6).

It is probably not surprising that some caudal *Hox* genes (i.e., *Hoxc6, Hoxc9*, and *Hoxd10*) that we observed to be up-regulated *in vitro* upon *Meg3* KD were not entirely phenocopied in the *IG-DMR^matΔ^* embryos, since this outcome is consistent with the finding that removal of *Ezh2* from MN progenitors has no detectable impact on segmental Hox expression in the spinal cord (Hoxc6 and Hoxc9) (Golden & Dasen, 2012). This result also suggests that a compensatory PRC1-mediated function *in vivo* might make up for the loss of *Meg3*-mediated epigenetic maintenance in segmental MNs. However, the significant Hoxa5-to-Hoxc8 homeotic transformation in MN subtypes supports the function of lncRNAs in mediating motor pool identity. Together with previous studies on stem cell-derived MNs and embryonic MNs, our results indicate that PRC2 plays dual roles at distinct phases in developing MNs. First, it establishes the chromatin landscape in the early pMNs. Then, together with the PRC1 complex, it consolidates segmental MN identity and maintains MN pool specification by decorating caudal Hox loci, particularly at the Hox8 region (Golden & Dasen, 2012; Mazzoni et al, 2013b).

### PRC2-lncRNA regulation in MNs

A previous study reported that hypomorphic *Suz12^−/−^* ESCs maintained with a low amount of H3K27me3 can differentiate into MNs, albeit with a significant increase in the expression of caudal *Hoxc6* and *Hoxa7* compared to wild-type cells (Mazzoni et al, 2013b). Interestingly, another study found that several PRC2 mutant ESC lines that maintain varying levels of H3K27me3 allowed for proper temporal activation of lineage genes during directed differentiation of ESCs to MNs, but only a subset of the genes that function to specify other lineages were not repressed in these cells (Thornton et al, 2014). This outcome might not be entirely surprising since other epigenetic marks, such as DNA methylation, might safeguard gene expression throughout differentiation (Manzo et al, 2017). In this study, up-regulated genes in spinal MNs upon loss of *Meg3-Rian-Mirg* exhibited 40% concordance (962/1598) with the up-regulated genes in *Suz12^−/−^* spinal MNs (data not shown). Altogether, these results strongly endorse the critical function of PRC2/lncRNA in perpetuating the postmitotic cell fate of cervical MNs.

Given that lncRNAs are also proposed to exert targeting of the PRC2 complex to specific genome loci (Rinn & Chang, 2012), it is tantalizing to hypothesize that *Meg3* might manifest dual functions of scaffolding the PRC2/Jarid2 complex and guiding it to specific loci in different cell contexts. This scenario could partially explain why only subsets of genes in an MN context are particularly sensitive to the loss of ncRNAs from the *Dlk1-Dio3* locus. Inspired by the salient Hox phenotype exhibited in the *IG-DMR^matΔ^* mutants, we envisage using *Meg3* as a paradigm to decipher the *Meg3*-protein-DNA interactome by ChIRP-seq/ChIRP-MS, thereby allowing us to decipher the detailed targeting mechanism of PRC2/Jarid2 involved in maintaining the epigenetic landscape during embryonic development (Chu et al, 2015).

### Combinatorial and individual roles of the *Meg3-Rian-Mirg* lncRNA cluster

There have been multiple efforts to dissect the functions of the maternally-expressed lncRNAs in the *Meg3-Rian-Mirg* locus over the past decade (McMurray & Schmidt, 2012; Steshina et al, 2006; Takahashi et al, 2009; Zhou et al, 2010), but definitive results remain elusive. The difficulty is mainly attributable to two major hurdles; namely that 1) there are many DMRs that control imprinting status in upstream, promoter, and exon regions of *Meg3*, and 2) *Meg3* might function *in cis* to regulate its imprinting status (Matsubara et al, 2015; Ogata & Kagami, 2016). Specifically, two *Meg3* knockout mouse lines have previously been generated either by deletion of the first five exons plus approximately 300 bp of the adjacent upstream promoter region of *Meg3* (~5.9 kb) (Yildirim et al, 2011) or by deletion of ~10 kb that includes the *Meg3-DMR* region plus the first five exons of *Meg3* (Takahashi et al, 2009). Both of these *Meg3* KO lines also manifested loss of maternal *Rian* and *Mirg* lncRNA expression. However, whereas the ~5.9 kb *Meg3* KO line (Yildirim et al, 2011) exhibited perinatal lethality, the ~10 kb *Meg3* KO line (Takahashi et al, 2009) presented a much milder phenotype in that the mice were born alive and lived up to 4 weeks after birth. It is important to point out that it is not possible to discern whether the effect of the *Meg3* deletion in either animal model is due to a lack of transcription from the *Meg3* promoter (leading to silencing of all lncRNAs), or due to a lack of *Meg3* expression that in turn causes silencing of downstream lncRNAs. We also found that expression of most, if not all, lncRNAs in the *Meg3*-*Rian*-*Mirg* locus are reduced upon *Meg3* KD and in *IG-DMR^matΔ^* mutants. Therefore, it remains technically challenging to obtain a specific *Meg3* KO without abrogating the expression levels of downstream lncRNAs.

Although we have further shown here that *Meg3* acts as a scaffold for the PRC2/Jarid2 complex, it is still possible that *Rian* and *Mirg* could independently and/or synergistically function with *Meg3* to contribute to the Hox-mediated MN subtype switching we observed in the *IG-DMR^matΔ^* mutants. Interestingly, *Rian* and *Mirg* KO ESCs/embryos seem to have a less severe phenotype and lack homeotic transformation (Han et al, 2014; Labialle et al, 2014). Thus, it is likely that *Meg3* is the major contributor to MN subtype specification in *IG-DMR^matΔ^* embryos. Since lncRNAs are emerging as important modulators of gene regulatory networks and as epigenetic regulators of gene expression, we are endeavoring to systematically knockout individual lncRNAs by a CRISPR-Cas9-mediated approach and we anticipate that a detailed map of lncRNA functions during neural development will be uncovered in the near future.

### Versatile functions of *Meg3* in development and disease

Consistent with the imprinting status of the *DLK1-DIO3* locus in humans, epimutations (hypermethylations) and microdeletions affecting *IG-DMR* and/or *MEG3-DMR* of maternal origin result in a unique human phenotype manifested as a small bell-shaped thorax, coat-hanger-like appearance of the ribs, abdominal wall defects, placentomegaly and polyhydramnios. One hallmark of patients with this disease, termed ‘Kagami-Ogata syndrome’ (KOS) (Kagami et al, 2015; Ogata & Kagami, 2016), is that nearly all of them display delayed gross motor development. It is currently unknown why epimutations and microdeletions of maternal *IG-DMR* give rise to this phenotype.

In our *IG-DMR^matΔ^* embryos, we previously observed extra ossification at the sites where the 6^th^ to 8^th^ ribs attach to the sternum, similar to the malformed thorax in KOS patients (Lin et al, 2007). Interestingly, this phenotype was also observed in *Hox5* mutant mouse embryos (McIntyre et al, 2007). Here, we uncovered that two isoforms of the *Meg3* imprinted lncRNA are enriched in embryonic MNs and confer the fidelity of the epigenetic landscape for the Hoxa5-Hoxc8 boundary of MN subtypes. Loss of *Meg3 in vitro* and *in vivo* abrogates the Hoxa5^on^ MNs in the brachial region, with a concomitant increase of ectopic Hoxc8^on^ subtypes. This switch leads to erosion of Hoxa5^on^ motor axon arborization in the proximal muscles, as well as eroded axon terminals along the tibialis anterior nerve. As *Meg3* expression is also highly enriched in somites, we suggest that impairment of the Hox boundary mediated by *Meg3* in the spinal cord and ribs might account for the bell-shaped thorax and motor deficit in KOS patients, potentially identifying a new therapeutic target for KOS patients.

In addition to the roles of the *Meg3* imprinted lncRNA uncovered by our study, other reports have also emphasized *Meg3* as being important for proper growth and development and to be a putative tumor suppressor that activates p53 and inhibits cell proliferation (Takahashi et al, 2009; Zhang et al, 2010; Zhou et al, 2007). Moreover, aberrant repression of *Meg3* and other maternally-expressed lncRNAs from the *DLK1-DIO3* imprinting cluster is present in several induced pluripotent stem cell (iPSC) lines. This scenario might lead to failure of these iPSCs to form viable mice (Stadtfeld et al, 2010) or to efficiently differentiate into neural lineage cells (Mo et al, 2015), raising the possibility that *Meg3* might be involved in a broad spectrum of developmental processes and disease contexts. Thus, our exploration of *Meg3* may also suggest new avenues for treating other diseases, such as cancers, as well as in elucidating the reprogramming mechanism of iPSCs.

## EXPERIMENTAL PROCEDURES

### Mouse ESC culture and MN differentiation

ESCs were cultured and differentiated into spinal MNs as previously described (Wichterle et al, 2002; Wichterle et al, 2009). Cells were trypsinized and collected for FACS at day 7 to purify GFP^on^ neurons for qPCR analysis and strand-specific RNA-seq when required.

### Mouse crosses and *in vivo* studies

The *IG-DMR^matΔ^* mouse strain is described in Lin et al. (2003). Female mice carrying the deletion were mated with wild type C57BL6/J male mice to generate embryos with the maternally-inherited deletion. Mice were mated at the age of 8~12 weeks and the embryo stage was estimated as E0.5 when a copulation plug was observed. Embryos were analyzed between E9.5~E13.5. All of the live animals were kept in an SPF animal facility, approved and overseen by IACUC Academia Sinica.

### Knockdown of *Meg3* in mouse ES cells by shRNA

The *Meg3* HuSH-29 shRNA plasmids (Origene®, cat. No. TG501330) and non-effective scrambled sequence (TR20003) were used to create stable knockdown lines of *Meg3* within the *Hb9::GFP* ESCs. We used two different shRNA sequences to knockdown *Meg3*. Additionally, stable infected ESCs were selected by puromycin. Single ESC clones with good morphology and only presenting knockdown efficiencies > 90 % were picked for further expansion and characterization.

### Expression analysis

ESCs or embryoid bodies were harvested for total RNA isolation by Trizol (Thermo Scientific). For qPCR analysis, total RNA from each sample was reverse transcribed with Superscript III (Thermo Scientific). One-tenth of the reverse transcription reaction was used for subsequent qPCR reactions, which were performed in triplicate with three independent experimental samples on a LightCycler480 Real Time PCR instrument (Roche) using SYBR Green PCR mix (Roche) for each gene of interest. *Gapdh* was used as a control for normalization. For GeneChip expression analysis, RNA was purified and amplified using the Qiagen RNAeasy kit and a one-color Low Input Quick Amp Labeling Kit (Agilent Genomics) and hybridized to a SurePrint G3 Mouse GE 8x60K Microarray. Differentially-expressed genes were defined by ranking all probes according to *Moderated t-test* and a fold-change threshold ≥ 2 (*P* < 0.001).

### Chromatin immunoprecipitation (ChIP)

We followed a previously published protocol to perform ChIP-seq for ESC~MNs (Mazzoni et al, 2011; Narendra et al, 2015). Four million cells were freshly dissociated from day 7 ESC~MNs by trypsin and fixed in 10 mM HEPES pH 7.6, 1 % formaldehyde, 15 mM NaCl, 0.15 mM EDTA and 0.075 mM EGTA for 15 minutes at room temperature. After fixation, cells were quenched with 1.25 M glycine. After an ice-cold PBS wash and low-speed centrifugation, nuclear extracts were suspended with ice-cold shearing buffer (SDS included) containing protease inhibitor and sheared using a Covaris M220 system to an average chromatin size of 200 bp. Chromatin was diluted with 2X IP buffer (2 % NP-40, 200 mM NaCl in 10 mM Tris-HCl pH 8, 1 mM EDTA). Anti-H3K27me3 antibody was added to each ChIP (antibodies are listed in the resource table). Each ChIP reaction was performed in a rotator at 4 °C overnight, followed by washing in wash buffer (25 mM HEPES pH 7.6, 1 mM EDTA, 0.1 % N-Lauryl sarcosine, 1 % NP-40, and 0.5 M LiCl) at room temperature. Cross-linking was reversed at 65 °C for 16 hours with 5 M NaCl. Proteinase K was added to digest for another 2 hours at 56 °C and DNA was extracted using the ChIP DNA Clean & Concentrator™ system (Zymo Research). We treated 1 % of the input in parallel. Libraries were prepared according to the Illumina protocol and sequenced using an Illumina NextSeq™ Sequencing System.

Whole-mount staining, immunohistochemistry, and *in situ* hybridization

Immunohistochemistry was performed on 20-µm cryostat sections as described (Chen et al, 2011). Primary antibodies used in this study are detailed in the resource table. Whole-mount antibody staining was performed as described (Dasen et al, 2008), and GFP-labeled motor axons were visualized in projections of a Zeiss Lightsheet Z.1 microscope (400–600 µm). Unless indicated otherwise, immunohistological data shown in figures are representative of n > 3 analyzed mutants. Images for control animals are from age-matched littermates. *In situ* hybridizations were performed as described previously (Chen et al, 2007; Chen et al, 2011) and in the Supplementary Experimental Procedures.

## ACKNOWLEDGMENTS

We thank people in the Chen lab, particularly to Hung Lo, for reading and giving critical comments to this manuscript. Dr. Mei-Yeh Lu from the NGS Core in Academia Sinica for invaluable technical advice and help in performing RNA-seq. Experiments and data analysis were performed in part through the use of the advanced optical microscopes at the Division of Instrument Services of Academia Sinica and with the assistance of Shu-Chen Shen. We appreciate the Genomic, FACS and Imaging core facilities in IMB for considerable technical assistance. The *IG-DMR^matΔ^* line was a kind gift from Prof. Ann Fergusson Smith of the University of Cambridge, UK, while the *Dbx1^Laz/+^* strain was from Prof. Alessandra Pierani from Institut Jacques Monod, CNRS UMR 7592, Université Paris Diderot, Sorbonne Paris Cité, France. The NIL plasmid was a gift from Prof. Hynek Wichterle from Columbia University, USA. We also acknowledge Bernhard Payer (CRG, Spain) and Wee-Wei Tee (A*STAR, Singapore) for their insightful and critical comments and for discussing the experimental results. The IMB’s Scientific English Editing Core reviewed the manuscript. This work is funded by Academia Sinica (AS-104-TP-B09), MoST (104–2311-B-001–030-MY3), and NHRI (NHRI-EX106–10315NC). SPL was supported by MoST (104–2321-B-002–043) and JHH was supported by MoST (104–2311-B-009–002-MY3 and 105–2221-E-009–126-MY3).

---

**Figure S1.**
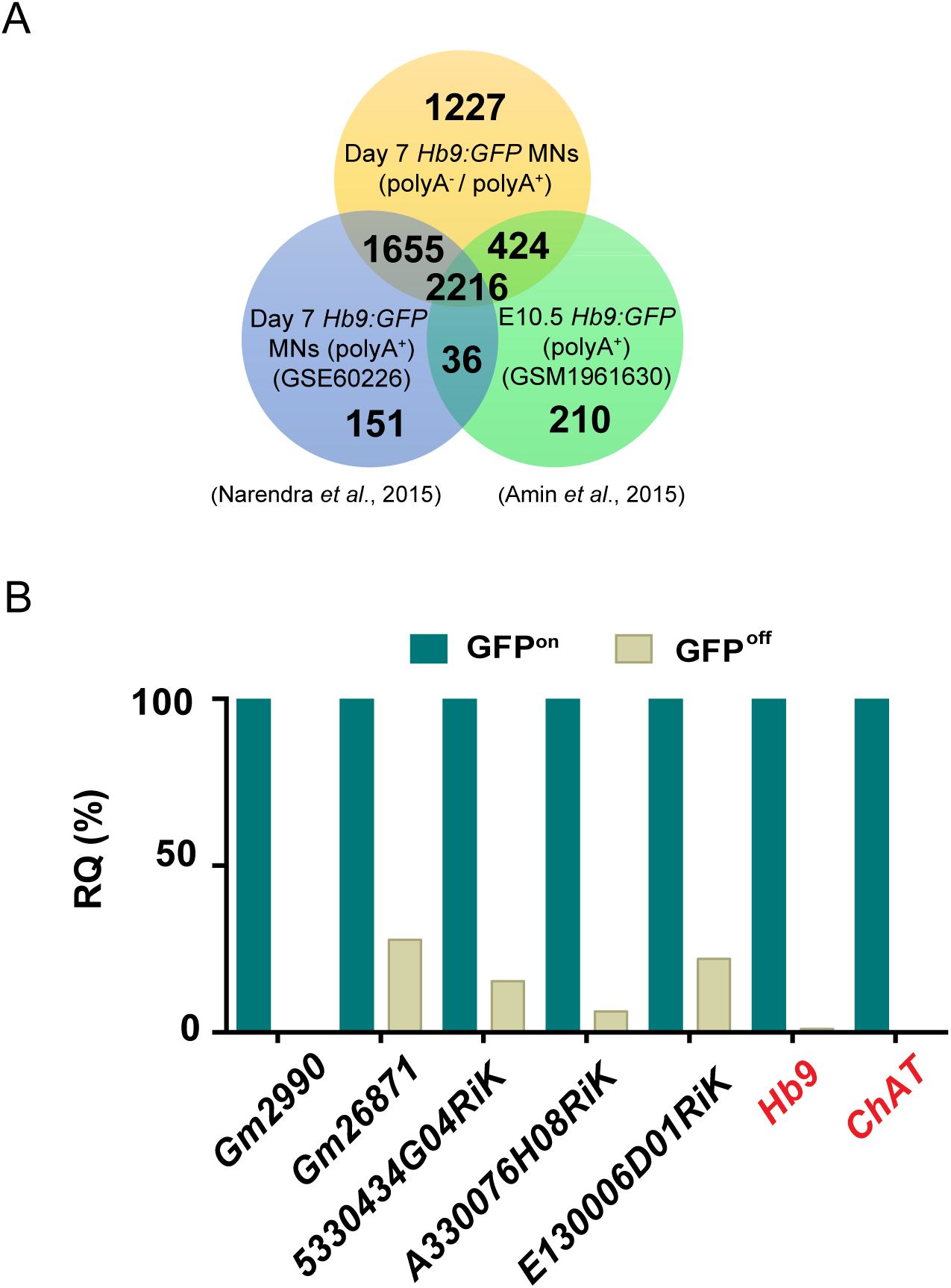
Systematic identification and verification of motor neuron signature lncRNAs. (A) The strand-specific RNA-seq performed in this study (orange) identified many more lncRNAs compared to previously published studies (blue and green) that used polyA^+^ selection RNA-seq. (B) Identified MN-signature lncRNAs were verified by qPCR from FACS-sorted GFP^on^ MNs, whereas GFP^off^ INs were used to reflect their specificity and relative quantity (RQ). *Hb9* and *ChAT* are mature MN markers.

**Figure S2.**
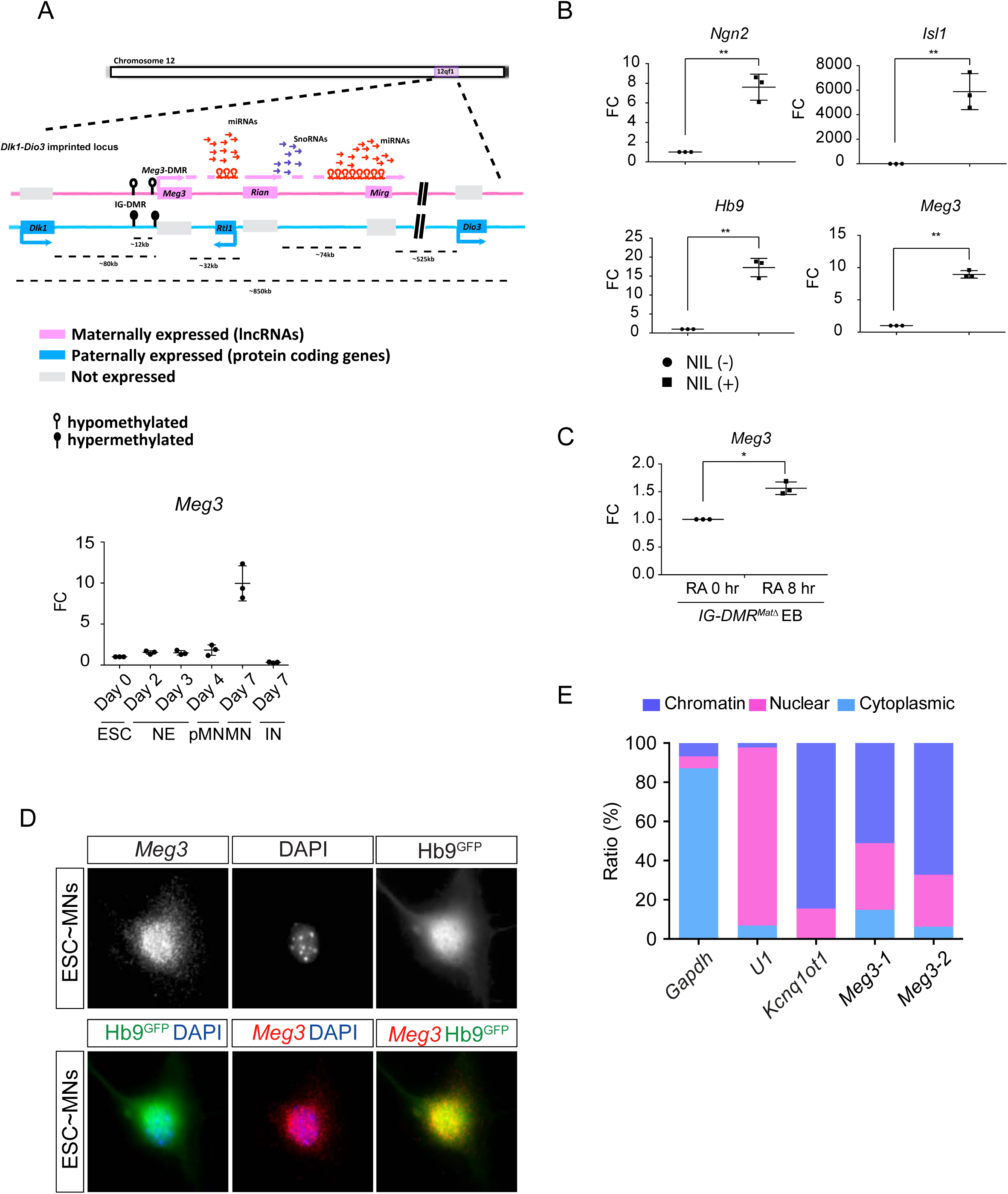
Characterization of the lncRNAs in the *Dlk1-Dio3* locus. (A) Illustration of the imprinted *Dlk1-Dio3* locus. The lncRNAs on the maternally-inherited allele (in pink) and protein-coding genes from the paternally-inherited allele (in blue) are depicted. miRNA genes are shown by hairpin loops. Arrows indicate the transcription directions. The *IG-DMR* site is hypomethylated (open circles) on the maternally-inherited allele, but hypermethylated (filled circles) on the paternally-inherited allele. Lower panel shows the qPCR analysis of *Meg3* expression in ESCs, NEs, pMNs, MNs and INs (n = 3 independent experiments; FC: fold-change). (B and C) qPCR reveals that MN transcription factors (*Ngn2-Isl1-Lhx3*), as well as adding RA, can induce *Meg3* expression in *IG-DMR^matΔ^* ESCs (FC: fold-change; error bars represent SD, n=3, ** p-value < 0.01, ** p-value < 0.05 by Student’s *t*-test). (D) Panels show single molecule RNA FISH of *Meg3* in *Hb9::GFP* ESC~MNs. (E) Subcellular fractionation of ESC~MNs revealed that *Meg3* is not only enriched in the nuclei, but also highly chromatin-associated. Two independent *Meg3* primers were used. *Gapdh*, *U1*, and *Kcnq1ot1* were used to reflect the purity of cytoplasmic, nuclear, and chromatin-associated fractions, respectively.

**Figure S3.**
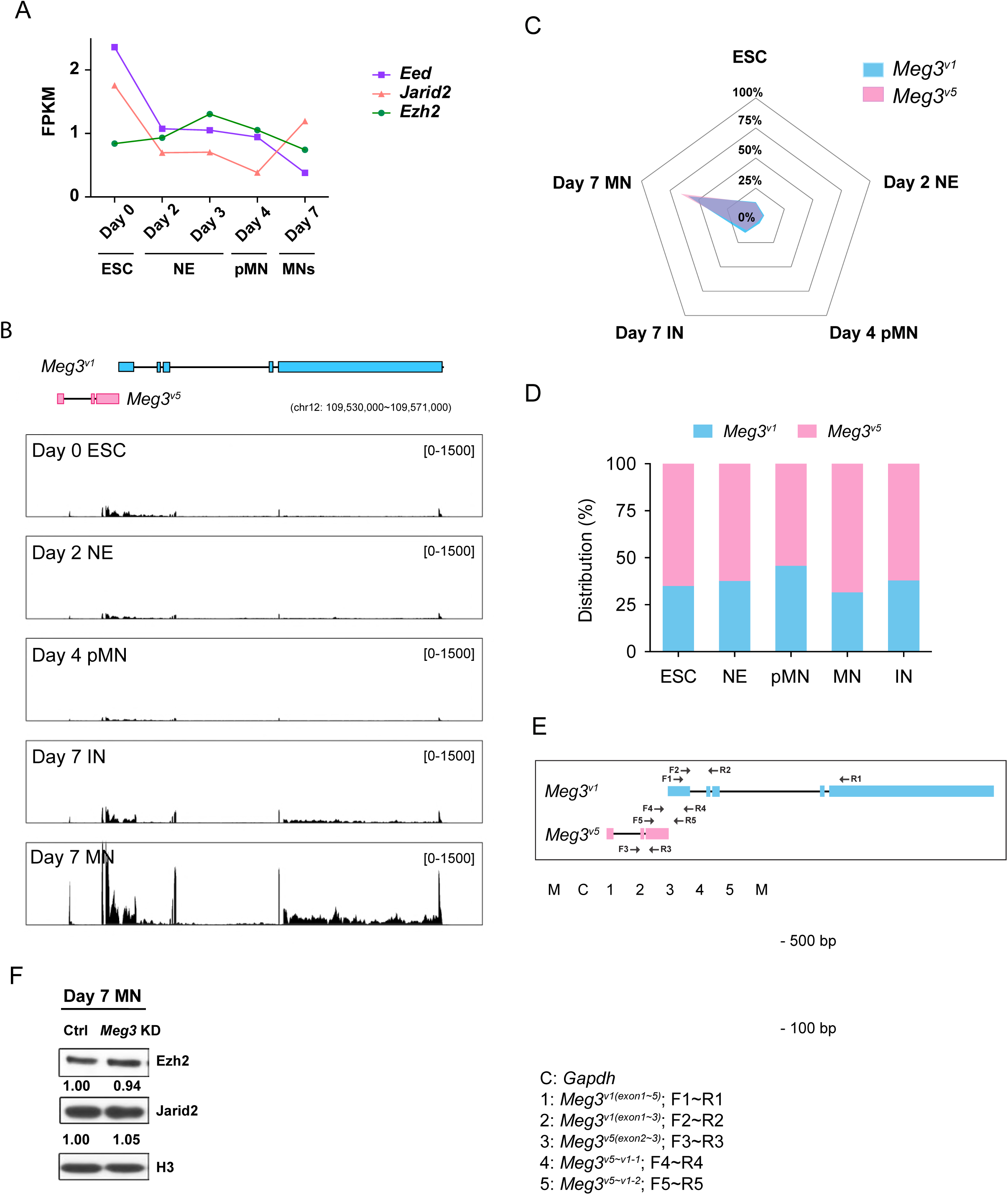
*Meg3* isoform characterization. (A) Time-series expression of the PRC2 subunits (Eed, Jarid2, and Ezh2) during ESC~MN differentiation. The levels of PRC2 complex progressively decreased during differentiation, whereas Jarid2 is reactivated in day 7 postmitotic MNs. (B) RNA-seq analysis of ESC~MNs. Reads from ESCs, RA-induced nascent neural epithelium (NE at day 2), MN progenitors (pMN at day 4), postmitotic interneurons (IN at day 7), and postmitotic MNs (MN at day 7). Reads are normalized to the total number of mappable reads per sample. RNA-seq revealed that *Meg3^v1^* (blue boxes) and *Meg3^v5^* (pink boxes) are the most abundant *Meg3* isoforms in postmitotic MNs (GENCODE version M9). (C) Radar chart revealing that day 7 postmitotic MNs have the highest distribution of *Meg3^v5^* during ESC~MN differentiation. (D) Histogram plot indicating that the *Meg3^v1^* and *Meg3^v5^* isoforms account for more than 99 % of *Meg3* transcripts during ESC~MN differentiation. (E) Schematic diagram of RT-PCR primer locations within the *Meg3^v1^* and *Meg3^v5^* regions. Expression of *Meg3* according to different primer combinations for *Meg3* isoforms suggests that *Meg3^v1^* and *Meg3^v5^* are independent transcripts. *Gapdh* as a loading control. (F) Western blot shows that the loss of *Meg3* imprinted lncRNAs does not affect the protein abundance of Ezh2 and Jarid2 in ESC~MN.

**Figure S4.**
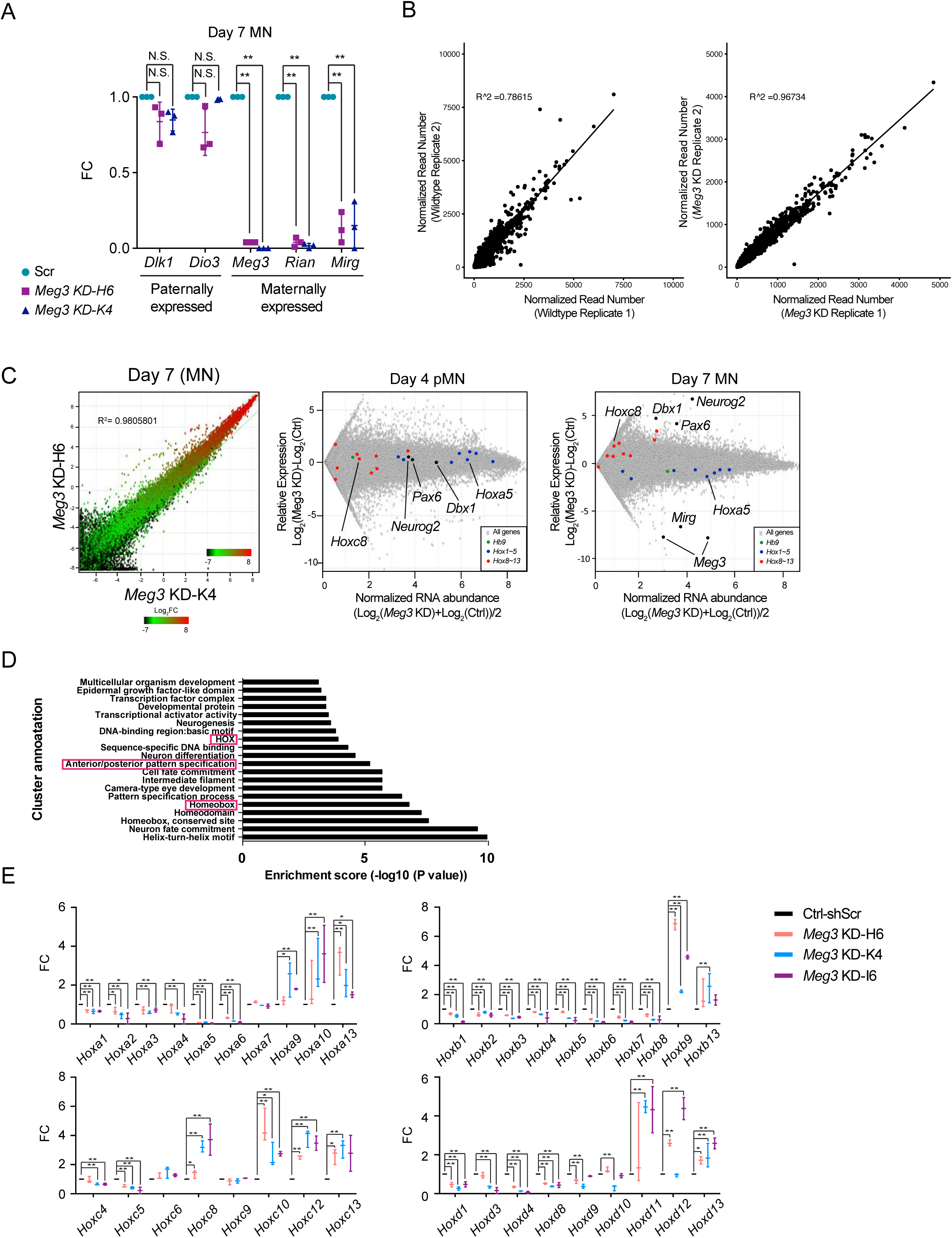
Characterization of *Meg3* KD ESCs. (A) Expression of paternally-expressed protein coding genes (*Dlk1* and *Dio3*) and maternally-expressed lncRNAs (*Meg3, Rian* and *Mirg*) in the imprinted *Dlk1-Dio3* locus, as revealed by qPCR (FC: fold-change; N.S.: not significant; error bars represent SD, n=3 independent experiments; * p-value < 0.05, ** p-value < 0.01 by Student’s *t*-test). (B) The correlation plots and associated R^2^ values are indicated by comparing H3K27me3 of MN samples from the two independent control and *Meg3* KD lines. (C) Right panel: The correlation plots and associated R^2^ values indicate that the two *Meg3* KD lines infected by independent shRNAs are almost identical. Left panel: MA plot of control versus *Meg3* KD pMNs (left) and MNs (right). At the postmitotic stage (day 7), *Meg3* knockdown leads to up-regulation of the neural progenitor genes *Pax6, Dbx1, Neurog2* and caudal *Hox* genes (*Hox8~13*), as well as down-regulation of rostral *Hox* genes (*Hox1~5*). Two biological replicate experiments were used to generate the data. The generic MN marker *Hb9* is unaffected, whereas *Meg3* and *Mirg* are the most down-regulated genes after *Meg3* KD. X-axis: mean abundance; Y-axis: log_2_ fold-change. (D) Dysregulated genes are grouped by gene ontology (GO). (E) Loss of *Meg3* leads to ectopic expression of most caudal *Hox* genes (*Hox8~13*), with concomitant down-regulation of rostral *Hox* genes (*Hox1~5*), as verified by qPCR. A third shRNA (I6) was further assessed and *Hox* expression was normalized against *Gapdh* expression levels (FC: fold-change; error bars represent SD, n=3; * p-value < 0.05, ** p-value < 0.01 by Student’s *t*-test).

**Figure S5.**
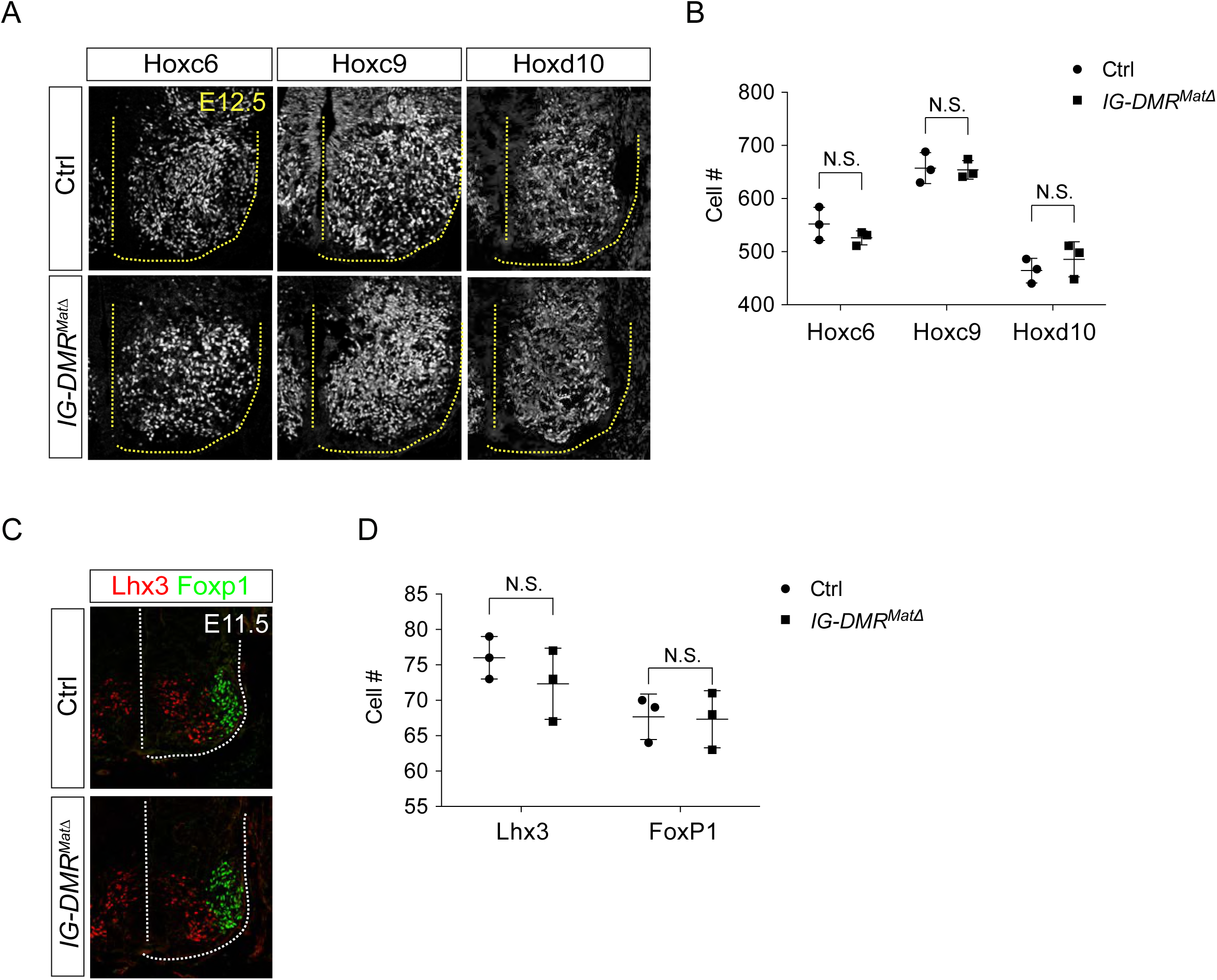
IG-*DMR^matΔ^* mutant phenotype analysis. (A and B) Immunostaining analysis of segmental MNs (Hoxc6^on^ at brachial, Hoxc9^on^ at thoracic, and Hoxd10^on^ at lumbar regions) show comparable MN numbers for E12.5 wild type control and *IG-DMR^matΔ^* spinal cord sections. Quantification of segmental MNs (number of positive cells per 15 µm brachial spinal cord ventral-half sections) in wild type control and *IG-DMR^matΔ^* embryos revealed no significant differences amongst different segments from the spinal cord (error bars represent SD, n=3 embryos at E12.5; N.S.: not significant). (C and D) Columnar axial MNs (Lhx3^on^) and limb-innervating MNs (Foxp1^on^) are unaffected in the *IG-DMR^matΔ^* embryos. Quantification of columnar MNs (number of positive cells per 15 µm brachial spinal cord ventral-half sections) in wild type control and *IG-DMR^matΔ^* embryos revealed no significant differences amongst different segments from the spinal cord (error bars represent SD, n=3 embryos at E11.5; N.S.: not significant).

**Figure S6.**
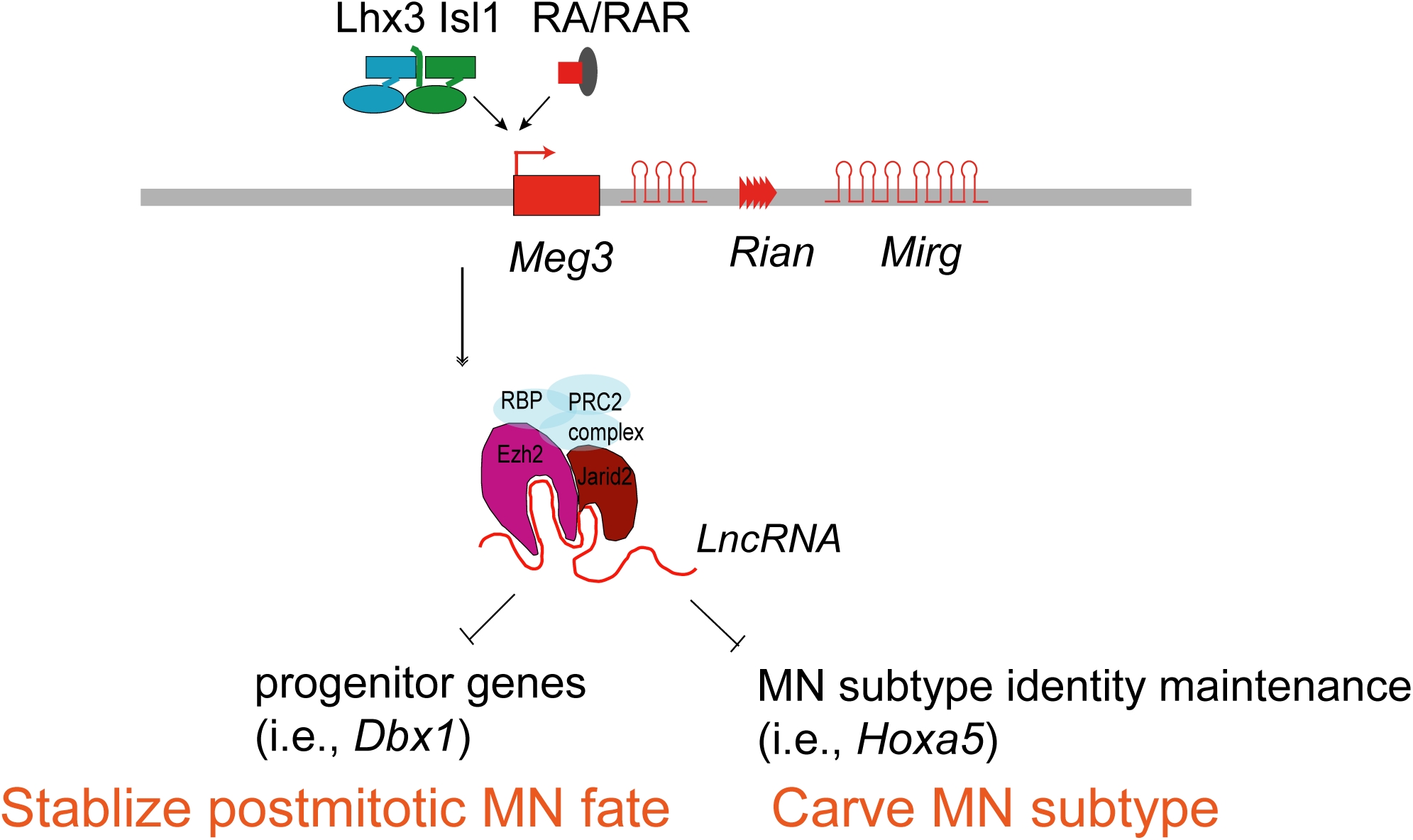
Summary of the functions of lncRNAs from the imprinted *Dlk1-Dio3* locus in MNs. *Meg3* and other lncRNAs from the *Dlk1-Dio3* locus are directly activated by MN-TFs (i.e Isl1 and Lhx3) and RAR, leading to enrichment of *Meg3* in the rostral segment of postmitotic MNs. One major function of *Meg3* and other lncRNAs from the *Dlk1-Dio3* locus is to stimulate Jarid2-Ezh2 interactions. Loss of these lncRNAs compromises the H3K27me3 epigenetic landscape and leads to aberrant expression of progenitor and caudal *Hox* genes in postmitotic MNs. Our model illustrates that the lncRNAs of the imprinted *Dlk1-Dio3* locus play a critical role in maintaining postmitotic MN cell fate by repressing progenitor genes, and that they shape MN subtype identity by regulating *Hox*.

## EXTENDED EXPERIMENTAL PROCEDURES

### KEY RESOURCES TABLE

| Reagent or Resource | Source | Identifier |
| --- | --- | --- |
| Rabbit polyclonal anti-Lhx3 | Abcam | ab14555 |
| Rabbit polyclonal anti-Foxp1 | Abcam | ab16645 |
| Goat polyclonal anti-Isl1 | Neuromics | GT15051 |
| Mouse monoclonal anti-Isl1(2) | DSHB | 39.4D5 |
| Rabbit polyclonal anti-Hoxa5 | Sigma-Aldrich | HPA029319 |
| Rabbit polyclonal anti-Hoxa5 | Jeremy Dasen (NYU) |  |
| Guinea pig polyclonal anti-Hoxa5 | Jun-An Chen (Academia Sinica) |  |
| Mouse monoclonal anti-Hoxc8 | DSHB | PCRP-HOXC8-1D3 |
| Rabbit polyclonal anti-Hoxc8 | Sigma-Aldrich | HPA028911 |
| Rabbit polyclonal anti-Pea3 | Thomas Jessell (Columbia University) |  |
| Guinea pig polyclonal anti-Scip | Thomas Jessell (Columbia University) |  |
| Sheep polyclonal anti-GFP | AbD Serotec | 4745-1051 |
| Mouse monoclonal anti-Nkx2.2 | DSHB | 74.5A5 |
| Rabbit polyclonal anti-Irx3 | Thomas Jessell (Columbia University) |  |
| Rabbit polyclonal anti-Olig2 | Millipore | AB9610 |
| Guinea pig polyclonal anti-Hb9 | Hynek Wichterle (Columbia University) |  |
| Mouse monoclonal anti-Hb9 | DSHB | 81.5C10 |
| Goat polyclonal anti-Hoxc6 | Santa Cruz | sc-46135 |
| Rabbit polyclonal anti-Hoxc9 | Jeremy Dasen (NYU) |  |
| Goat polyclonal anti-Hoxd10 | Santa Cruz | sc-33005 |
| **RNA/Protein Immunoprecipitation (RIP/IP) and Western blot** |  |  |
| Reagent or Resource | Source | Identifier |
| Mouse monoclonal anti-Ezh2 | Millipore | AC22, 17-662 |
| Rabbit polyclonal anti-Jarid2 | Novus | NB100-2214 |
| Rabbit polyclonal anti-Suz12 | Abcam | ab12073 |
| Mouse monoclonal anti-H3K27me3 | Abcam | ab6002 |
| **Chromatin Immunoprecipitation (ChIP)** |  |  |
| Reagent or Resource | Source | Identifier |
| Mouse monoclonal anti-H3K27me3 | Abcam | ab6002 |
| **Experimental Models: Cell Lines** |  |  |
| Reagent or Resource | Source | Identifier |
| Mouse: *Hb9::GFP* ESC | (Wichterle et al. 2002) |  |

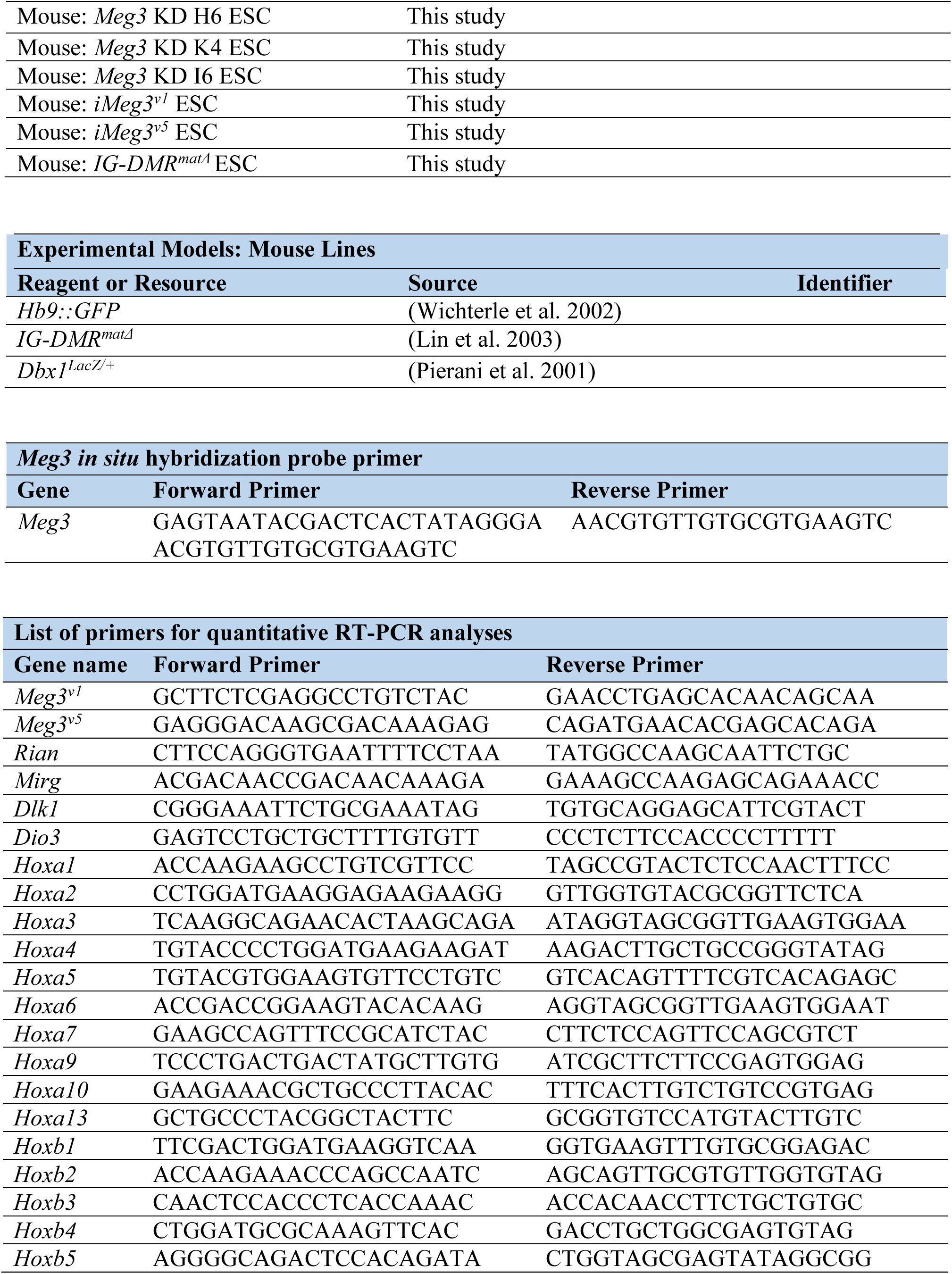

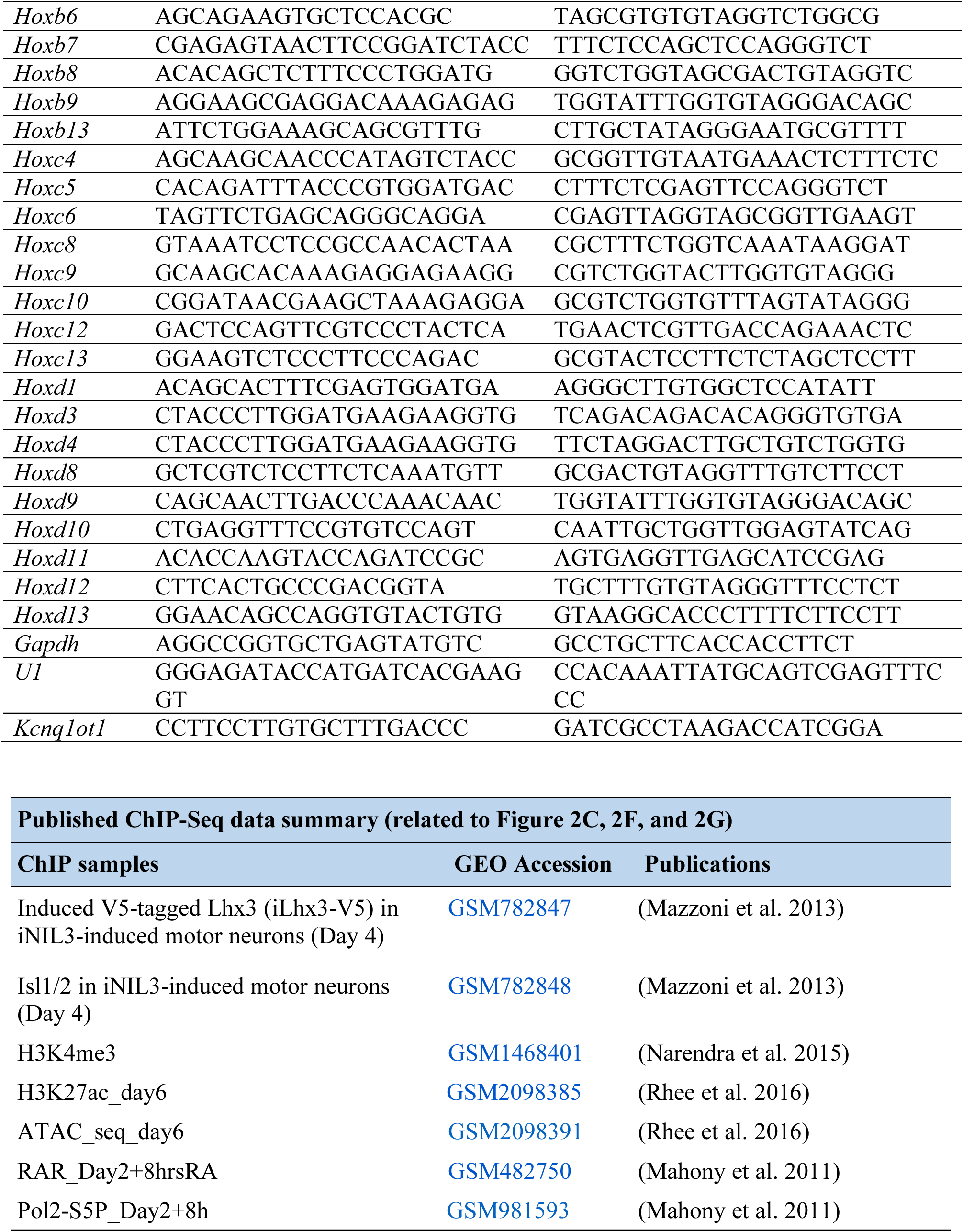

Further information and requests for reagents may be directed to, and will be fulfilled by, the Lead Contact, Jun-An Chen.

## METHOD DETAILS

### Immunocytochemistry

Embryos and/or embryoid bodies were fixed with 4 % paraformaldehyde (vol/vol) in phosphate-buffered saline, embedded in OCT (Tissue-Tek) and sectioned for staining; 24 hours at 4 °C for primary antibodies and 1~2 hours at 20~25 °C for secondary antibodies. Alexa488-, Cy3- and Cy5-conjugated secondary antibodies were obtained from either Invitrogen or Jackson Immunoresearch. After staining, samples were mounted with Fluoroshield with DAPI (Roche). Images were acquired with a Zeiss LSM 710/780 or LightSheet Z. 1 confocal microscope.

### Subcellular RNA fractionation

We followed a previously published protocol to extract subcellular fractions of RNA (Gagnon et al. 2014). We used TRIzol (Thermo Fisher Scientific) to extract RNA and perform reverse transcription (RT) with hexamer primers. *Gapdh* (mRNA in cytoplasm), *U1* (snRNA in nucleus), and *Kcnq1ot1* (a known chromatin-associated lncRNA) were used as quality controls to verify fractionation.

### RNA pull-down assay

In *vitro*-transcribed biotin-labelled RNAs were generated by the Biotin RNA Labeling Mix (Roche) and T7 RNA polymerase (Promega). Templates were treated with RNase-free DNase I (Promega) and the reaction mix was purified with Oligo Clean & Concentrator™ (D4060, Zymo Research). Biotinylated RNA (3 µg) was heated to 65 °C for 10 minutes and then slowly cooled down to 4 °C. After that, RNA structure buffer (10 mM Tris pH 7, 0.1 M KCl, 10 mM MgCl_2_) was added and the mix was shifted to room temperature for 20 minutes to allow proper secondary structure formation. Folded RNA was then mixed with 1 mg of ESC protein nuclear extract in RIP buffer (500 mM NaCl, 10 mM HEPES pH 7.5, 25 % glycerol, 1 mM EDTA, and protease inhibitor) and incubated at 4 °C for one hour. Twenty µL Dynabeads® M-280 Streptavidin (Invitrogen) were added to each binding reaction and further incubated at room temperature for one hour. Beads were washed briefly five times and boiled in SDS buffer, and the retrieved protein was detected by standard Western blot analysis.

### Co-immunoprecipitation (Co-IP) and Western blot

For each IP, cells were harvested from a 10-cm dish and washed twice with ice cold PBS. Cell pellets were allowed to swell in twice the volume of cytoplasmic lysis buffer (50 mM NaCl, 10 mM HEPES-pH 7.5, 500 mM sucrose, 1 mM EDTA and protease inhibitors). Samples were incubated on ice for 10 minutes, followed by centrifugation at 2,000 rpm for 10 minutes. The cloudy supernatant cytoplasmic fraction was removed. After washing twice (50 mM NaCl, 10 mM HEPES-pH 7.5, 25 % glycerol, 1 mM EDTA and protease inhibitors), the cell pellets were resuspended in the same volume of high salt buffer (500 mM NaCl, 10 mM HEPES-pH 7.5, 25 % glycerol, 1 mM EDTA and protease inhibitors), and rotated for 30~60 minutes at 4 °C. Then cell pellets were centrifuged at 14,000 rpm for 10 minutes at 4 °C. The supernatant was incubated overnight at 4 °C with antibody and pre-cleared Protein-G beads (depending upon the antibody) to immunoprecipitate endogenous protein against the specific antibody used. We collected 10 % of cleared supernatant as input. Subsequently, IP-protein beads were washed three times with PBS and 0.01 % Tween-20, each for 5 minutes at 4 °C. IP-proteins and their interacting partners were eluted from beads in 6X reducing loading buffer at 70 °C for 15 minutes. Finally, samples were cooled down to room temperature and spun briefly to collect condensation. Standard Western blot procedures were applied using anti-Jarid2 (Novus Biologicals, NB100–2214) or anti-Ezh2 (Millipore, 17–662) antibodies. Blots were developed using HRP-conjugated anti-rabbit or -mouse antibodies, depending on the species of the primary antibody. Signals were developed and filmed by enhanced SuperSignal™ West Femto Maximum Sensitivity Substrate (Thermo, 34096). All exposures were done using hyper film.

### Single molecular RNA FISH

ESC~MNs were cultured and harvested on slides. Cells were fixed in 4 % paraformaldehyde for 10 minutes at room temperature, permeabilized for 5 minutes on ice in PBS with 0.5 % Triton X-100, and then rinsed in 70 % EtOH for subsequent RNA FISH. Slides and coverslips were kept in 70 % EtOH at 4 °C until staining. Slides were then washed in wash buffer (10 % deionized formamide in 2X SSC) for 5 minutes and incubated in a dark room at 37 °C for at least 4 hours with 1 µL of probe stock solution and 100 µL of hybridization buffer (1 g dextran sulfate, 1 mL 20X SSC, 1 mL deionized formamide). *Meg3* smFISH probes were purchased from Stellaris. Images were captured with a Delta Vision microscopy system.

### RNA immunoprecipitation (RIP)

RIP was performed with the RNA-Binding Protein Immunoprecipitation Kit (17–700, Millipore) according to the manufacturer’s protocol with some modifications. ESC~MNs were dissociated at a concentration of 2 million cells/mL and treated with 0.3 % formaldehyde in ice-cold PBS for 10 minutes at 37 °C. Glycine/PBS was added to a final concentration of 0.125 M and each sample was incubated for 5 minutes at room temperature. After crosslinking, ten million cells were washed twice with cold PBS and then suspended in 100 µL RIP lysis buffer (with the addition of protease inhibitor and RNase inhibitor). The lysate was incubated on ice for 5 minutes and centrifuged at 14,000 rpm for 10 minutes at 4 °C. Ezh2, Jarid2, and Suz12 antibodies were added for respective IP reactions and then incubated in RIP buffer (0.5 M EDTA/RNase inhibitor) for 3 hours to overnight at 4 °C. Samples were washed at least five times with RIP washing buffer. RIP beads were resuspended in RIPA buffer (RIP washing buffer + 10 % SDS + protease K) to reverse crosslinking at 56 °C for 30 minutes. RNA samples were extracted and qPCR was performed as described above. Isolated proteins before proteinase K treatment were collected from the beads and verified by Western blot analysis. Data on retrieved RNAs was calculated from the RT/input ratio for each experiment.

### Statistical analyses and graphical representations

All statistical analyses were generated with GraphPad Prism 6 (GraphPad Software). The values are shown as mean ± SD, as indicated. Student’s *t*-tests were used for comparisons between experimental samples and controls. Statistical significance was defined as *p < 0.05 and **p < 0.01 by Student’s *t*-test.

### Bioinformatics procedures

#### RNA-seq analysis

Adapter contamination in the paired-end reads was removed using PEAT (Li et al. 2015), and the trimmed reads were aligned to the mm10 genome with STAR (Dobin et al. 2013). The standard GTF-formatted transcript annotation was defined by GENCODE (version M9) (Harrow et al. 2012), which includes many evidence-based lncRNAs. We used this annotation to aid the junction read alignment in STAR, the output of which was submitted to Cufflinks (Trapnell et al. 2010) for *de novo* transcript assembly with the option ‘library type; first-strand’ to allow strand-specific alignments. We followed a strategy for novel lncRNA identification similar to that suggested by a previous report (Qian et al. 2016), by which only transcripts that were longer than 200 bp, had no overlap with any known genes, and consisted of more than one exon were regarded as novel lncRNAs. We pooled these novel lncRNAs along with all known genes annotated in GENCODE and used HTseq (Anders et al. 2015) to calculate the read count aligned onto each transcript. This procedure was repeated for all RNA-seq samples in this study. The read counts of all transcripts among different samples were normalized using a TMM algorithm with the trimming option M=30 % and A=5 % (Robinson and Oshlack 2010). A general comparison of different normalization algorithms can be found in Lin et al (Lin et al. 2016). We calculated the specificity score of each transcript among the samples at different stages according to the Jensen–Shannon definition for tissue specificity scores (Trapnell et al. 2010; Cabili et al. 2011). The transcripts were split into three groups—namely protein coding genes, annotated lncRNAs, and novel lncRNAs—for which specificity score distributions were plotted and compared.

#### ChIP-seq analysis

Reads were trimmed by PEAT and aligned to the mm10 genome using Bowtie2 (Langmead and Salzberg 2012). Following a similar flow analysis described in our previous work (Yildirim et al. 2011; Chen et al. 2013), all alignments were extended downstream to span an exact 150 bp-long region. Extensions that exceeded the ends of chromosomes were clipped. The extended alignments were input into the *genomecov* functionality supported in the BEDTools suite (Quinlan and Hall 2010) to generate read coverage profiles at a base-pair resolution. The coverage for each chromosomal position was normalized according to the mappable read count. Each sample was averaged and binned to reveal major trends. To identify possible differentially-enriched histone marks among stages or treatments, we used MACS 1.4 (Feng et al. 2011) to call peaks (P-value <10^−5^) in each ChIP-seq sample with the corresponding input library and then overlapping peaks were merged using MAnorm (Shao et al. 2012) to reveal loci with a significant change between two samples.

#### Axon arborization quantification with Imaris

The 3D images acquired with a Zeiss Lightsheet Z.1 microscope were subjected to analyses in Imaris 8.4.0 (Bitplane, Zurich, Switzerland) for quantification of axon arborization. Regions of interest were segmented for detection of individual neurons. Motor nerve terminals were semi-automatically traced using the filament tracer wizard from a defined starting point. The AutoPath (no loops) algorithm was selected. Seed points detected from background signals were manually removed. Disconnected segments were removed by indicating the maximum gap length, and background subtraction was applied for noise removal. The “Filament No. Dendrite Terminal Points” tool automatically calculated the number of motor nerve terminals.

